# Brain-thought states during movie-watching reflect shared sensory anchors and film- and person- specific processes in association cortex

**DOI:** 10.64898/2026.10.16.756266

**Authors:** Raven S. Wallace, Samyogita Hardikar, Samuel Ketcheson, Bridget Mulholland, Ian Goodall-Halliwell, Bronte McKeown, Louis Chitiz, Michael Milham, Ting Xu, Daniel S. Margulies, Nerissa Ho, Theodoros Karapanagiotidis, Giulia L. Poerio, Tirso R. J. Gonzalez Alam, Adam Turnbull, Jeremy I. Skipper, Boris C. Bernhardt, Robert Leech, Jeffrey D. Wammes, Elizabeth Jefferies, Jonathan Smallwood

## Abstract

Naturalistic paradigms such as movie-watching have transformed functional neuroimaging by capturing cognitive dynamics during rich, real-world experiences. A fundamental challenge, however, is determining whether the neural signatures identified during these experiences reflect general principles of cognition or the specific contexts in which they are measured. To address this question, we combined fMRI data from participants watching eight diverse films with continuous reports of ongoing thought. We found that the generalizability of brain-thought relationships depends on the cognitive process being recruited. Neural signatures linked to sensory processing remained robust across films, whereas those involving memory-related association networks depended on how we modelled between-film variability, indicating that cognition becomes increasingly context-dependent along the cortical hierarchy from sensory to association cortex. Within default and limbic networks, we identified two contrasting memory-related states with opposite consequences for comprehension. One, reflecting the integration of prior knowledge with ongoing experience, predicted better narrative comprehension and was revealed only after accounting for between-film variability. The other, associated with stimulus-independent thought, predicted poorer comprehension and emerged only when between-film variability was not modelled. Together, these findings establish when brain-experience relationships measured during naturalistic paradigms can generalize across contexts and when they instead reflect interactions between cognition and the specific environments in which cognition unfolds. More broadly, the consistency of a given brain-experiential association may depend less on where it occurs in the brain and more on the viewer’s contextual experience: perceiving the film, drawing on memory to understand it, or drifting away.

**Significance statement:** Films have revolutionized how we study healthy and atypical brain activity by capturing synchronized, semi-unconstrained dynamics of real-world cognition. Yet experience is rarely measured alongside brain activity, and most studies use one or few films; thus, we do not know whether these neural and experiential responses generalize across films. By combining brain imaging with continuous reports of ongoing thought, we find a hierarchy: sensory processing is consistent across films, whereas memory-related processes in association cortex are uniquely sensitive to how film context is modelled. Two opposing memory states relate to comprehension. One draws on memory to support understanding; the other reflects stimulus-independent thought and impairs it. Identifying brain–experience relationships across films is crucial for developing and understanding naturalistic neuroscience.

## 1. Introduction

Naturalistic paradigms, such as movie-watching, have become a central tool in neuroimaging, especially in functional magnetic resonance imaging (fMRI). Unlike traditional laboratory tasks, which isolate specific cognitive components (Brewer et al., 1998; Cabeza & Nyberg, 2000; Kanwisher et al., 1997; MacDonald et al., 2000; Owen et al., 2005), film paradigms evoke perception, emotion, and meaning that resemble the spontaneous, overlapping processes of real-world experience (Finn, 2021; Kringelbach et al., 2023). Conventional resting-state paradigms have led to important characterizations of intrinsic functional connectivity networks. However, their underlying interpretation remains inherently ambiguous given the breadth of cognitive states one could occupy during a task-free paradigm (Biswal et al., 1995; Cole et al., 2010; Fox & Raichle, 2007; Gusnard et al., 2001; Smitha et al., 2017). Features of brain function during movie-watching better predict individual differences in cognition and emotion than those captured during resting state and have increasingly been used in developmental and clinical research (Eickhoff et al., 2020; Finn, 2021; Finn & Bandettini, 2021; Gal et al., 2022; Samara et al., 2023; Vanderwal et al., 2017).

Neural responses to the same film are often strikingly similar across individuals (Hasson et al., 2008a; Hasson et al., 2004), and intersubject correlation (ISC) and event boundary research show substantial alignment in how viewers segment and encode films (Barnett et al., 2024; Hasson et al., 2008b; Zacks et al., 2001). These commonalities make movie stimuli useful for inducing particular shared cognitive states (Yeshurun et al., 2021). However, this consistency is typically established across viewers of the same film. Many studies rely on one or a few films, implicitly treating the stimulus as a fixed design feature rather than a sample from a larger population of films (Yarkoni, 2020). Films themselves differ substantially in narrative, genre, cultural origin, mood, and reliance on dialogue (Leipold et al., 2025), all of which influence neural responses. Thus, when we identify relationships between brain activity and experience, we need to discern whether they reflect a general feature of cognition or a feature of the particular film measured.

Whether brain-experiential relationships generalize across films may depend on where they sit along the cortical hierarchy, as movie-watching dynamically recruits both lower-level sensory cortices and higher-order association regions (Hasson et al., 2008a; Song et al., 2021b; Wallace et al., 2025). Sensory regions have relatively well-defined roles associated with specific perceptual modalities: they track changes in the films’ physical properties on shorter timescales and respond consistently across viewers (Hasson et al., 2004; Hasson et al., 2008b; Samara et al., 2023). Heteromodal association systems, particularly the default mode network (DMN), integrate information over longer timescales and sit at the apex of the principal gradient of cortical organization extending from sensory-motor to association cortex (Margulies et al., 2016), and their large-scale functional topography shifts with cognitive states (Wang et al., 2025). Unlike sensory cortex, the recruitment of these regions depends on what a film contains: neuroimaging during a non-social, non-verbal movie paradigm (e.g., *Inscapes*) resembles resting state more than conventional narrative films (Vanderwal et al., 2015), and films that vary in social and narrative content differentially engage cortex beyond sensory regions (Vanderwal et al., 2019; Vanderwal et al., 2017). Consistent with these findings, association regions are prominently engaged by evolving narratives (Cooper et al., 2021; Yang et al., 2023), but they also show the greatest variation in neural responses across individuals and films (Kröll et al., 2023; Leipold et al., 2025; Song & Wang, 2024). *If association cortex supports processes that depend on how a particular narrative interacts with what a viewer brings to it, brain-experiential relationships should become increasingly context-dependent moving from sensory to association cortex*.

Nonetheless, interpreting associations with higher-order heteromodal systems can be complex. Association regions support a wide range of cognitive states, from comprehending meaningful visual and verbal inputs to mind-wandering that can disrupt task performance (Andrews-Hanna et al., 2010; Christoff et al., 2009; Paquola et al., 2022; Paquola et al., 2025; Poerio et al., 2017; Smallwood et al., 2021a; Yeshurun et al., 2021). During movie-watching, DMN activity has been linked to processes that support understanding, such as segmenting events, retrieving relevant knowledge, and integrating information into a coherent narrative (Baldassano et al., 2017; Lee & Chen, 2022; Song et al., 2021a). Yet the same regions are implicated in mind-wandering and perceptual decoupling, states that disrupt comprehension of ongoing input (Christoff et al., 2009; Poerio et al., 2017; Smallwood, 2011; Zhang et al., 2022). During films, DMN regions also covary with changes in attention and affect, reflecting moments of synchronization and engagement during emotionally charged and surprising events, as well as moments of distraction and disengagement (Bezdek et al., 2017; Jacob et al., 2018; Nanni-Zepeda et al., 2024; Song et al., 2021a). Together, these findings suggest that the contribution of higher-order association cortex during movie-watching is more difficult to characterize than that of sensory systems because it supports multiple, potentially competing, cognitive processes. Functional neural data alone cannot distinguish these states, as increased activation of DMN regions could reflect both deep engagement with the film and a departure from its content (Jefferies & Smallwood, 2025; Yeshurun et al., 2021). Thus, resolving this ambiguity requires integrating the viewer’s experience as the film evolves with behavioural measures of comprehension to understand which states support or impair memory for these films.

Differentiating whether shared mechanisms truly support movie-watching cognition requires an analytic approach that treats films as a source of variance in their own right; just as participants are modelled as random effects, films can be modelled as random effects to test whether conclusions generalize beyond the sampled stimuli to the broader population of films (Yarkoni, 2020). Accounting for between-film variance can alter brain-experiential relationships in opposing ways. On the one hand, it can reveal relationships that hold within films but are masked by systematic differences between them; on the other, it can eliminate relationships that exist only because films differ in the brain and experiential states they evoke. Thus, comparing models that do and do not account for film identity can provide a principled approach to determining which brain-experiential relationships are stimulus-invariant and which depend on the particular films sampled.

Here, we mapped ongoing brain activity onto the multifaceted experience of movie-watching across eight films using openly available fMRI data from the Naturalistic Neuroimaging Database (NNDb; n = 73; Aliko et al., 2020). To capture participants’ subjective experiences, an independent sample (*n* = 226) watched clips from the same films while completing multi-dimensional experience sampling (mDES; see (Smallwood & Schooler, 2015; Smallwood et al., 2021b; Wallace et al., 2025)), where participants periodically report on multiple aspects of their experience, such as level of detail, temporal orientation, focus, and reliance on different sensory modalities. To avoid disrupting the naturalistic viewing experience, we used an optimized sampling regime from prior work to minimize the number of probes each participant receives (five per clip) while maintaining dense coverage aggregated across the sample, yielding experiential reports every 10–15 seconds. After each clip, participants additionally completed a brief comprehension test on the film’s content. To situate brain activity within a coordinate system independent of any single film, we projected moment-to-moment activity onto the first five gradients of intrinsic cortical organization derived from resting-state data (Margulies et al., 2016; Wallace et al., 2025). Extending random-effects modelling beyond participants to stimuli (Yarkoni, 2020), we compared a baseline model with participants as random intercepts with a crossed model that also accounted for film-level variance. We hypothesized that brain–experience relationships anchored in sensory systems would be robust to between-film variability, whereas those involving association cortex would be most affected by modelling film-level variance, reflecting these regions’ greater functional variability and their high sensitivity to unique contextual features during movie-watching.

## Results

### 2. Analytic Goal

We linked the thought patterns experienced across diverse movies with brain activity using two separate samples who watched the same films: one that provided experiential annotations during movie-watching using mDES (*n* = 226; 185 women, *M+SD*_age_ = 19.03±1.30 years; range = 17-30 years), and another, from the Naturalistic Neuroimaging Database (NNDb, (Aliko et al., 2020)) that had watched the films during uninterrupted fMRI sessions (*n* = 73; 35 females, *M±SD*_age_ = 27.63±10.82 years, range = 18-58). The analysis was performed using 9- to 11-minute clips from eight different movies, including *Citizenfour* (Documentary)*, Little Miss Sunshine* (Comedy)*, 500 Days of Summer* (Romance)*, Back to the Future* (Sci-Fi*, Shawshank Redemption* (Drama)*, 12 Years a Slave* (Historical)*, The Prestige* (Thriller), and *Pulp Fiction* (Action). The mDES probes assessed thoughts across 16 dimensions describing features of ongoing experience, including emotional valence, temporal dynamics, and sensory features (for example, “My thoughts were in the form of images” or “My thoughts involved myself”; for the complete questionnaire, see **Supplementary Table 12**). In this study, we use mDES probes administered during movie-watching to establish thought patterns and to assess their association with corresponding brain activation patterns across multiple film genres.

### 3. Generation of thought and brain spaces

Our approach involves creating low-dimensional ‘thought’ and ‘brain’ spaces by reducing our 16-item mDES data and voxel-dimension brain data to show how patterns of experiential reports and brain activity emerge across our selected films. First, following prior work, we used Eigenvector decomposition, to map the mDES data reported by participants across all films into a set of ‘thought dimensions’ (Konu et al., 2021; McKeown et al., 2025; McKeown et al., 2021; McKeown et al., 2023; Mulholland et al., 2025; Mulholland et al., 2023; Wallace et al., 2025; Zhou et al., 2025) (see **Figure 1**, bottom right). We extracted a five-component solution that explained 56.6% of the total variance, in which the loadings of each retained component describe the features of experience. The loadings for each thought pattern can then be visualized with word clouds, where red and blue represent positive and negative loadings, respectively; similar colours are more closely related, and the font size indicates the magnitude of the loading, an indication of the extent to which that dimension drives a given thought pattern (see **Figure 1**). For simplicity, going forward, we assigned each component a descriptive label based on the items with the most prominent loadings. For instance, Component 1 loaded positively on “distracting” and “intrusive” and negatively on “deliberate” and was labelled *Intrusive Distraction*, while Component 2 loaded positively on “sounds,” “images,” and “people” and is referred to as *Sensory Engagement*. Components 3, 4 and 5 were labelled *Episodic Knowledge*, *Deliberate Detail,* and *Positive Engagement,* respectively (See **Supplementary Table 1** for the rotated component matrix). While these specific thought patterns were derived within this dataset, they align nearly perfectly with those observed in many previous samples in the lab (Konu et al., 2021; McKeown et al., 2025; Turnbull et al., 2021; Wallace et al., 2025; Zhou et al., 2025), daily life (McKeown et al., 2021; McKeown et al., 2023; Mulholland et al., 2025; Mulholland et al., 2023), and even when accompanied by changes in brain activity during fMRI scanning (Konu et al., 2020; Turnbull et al., 2019). Because each probe response can be described using scores on each of the five retained components, these thought patterns create a 5-dimensional ‘thought space’.

**Figure 1.**
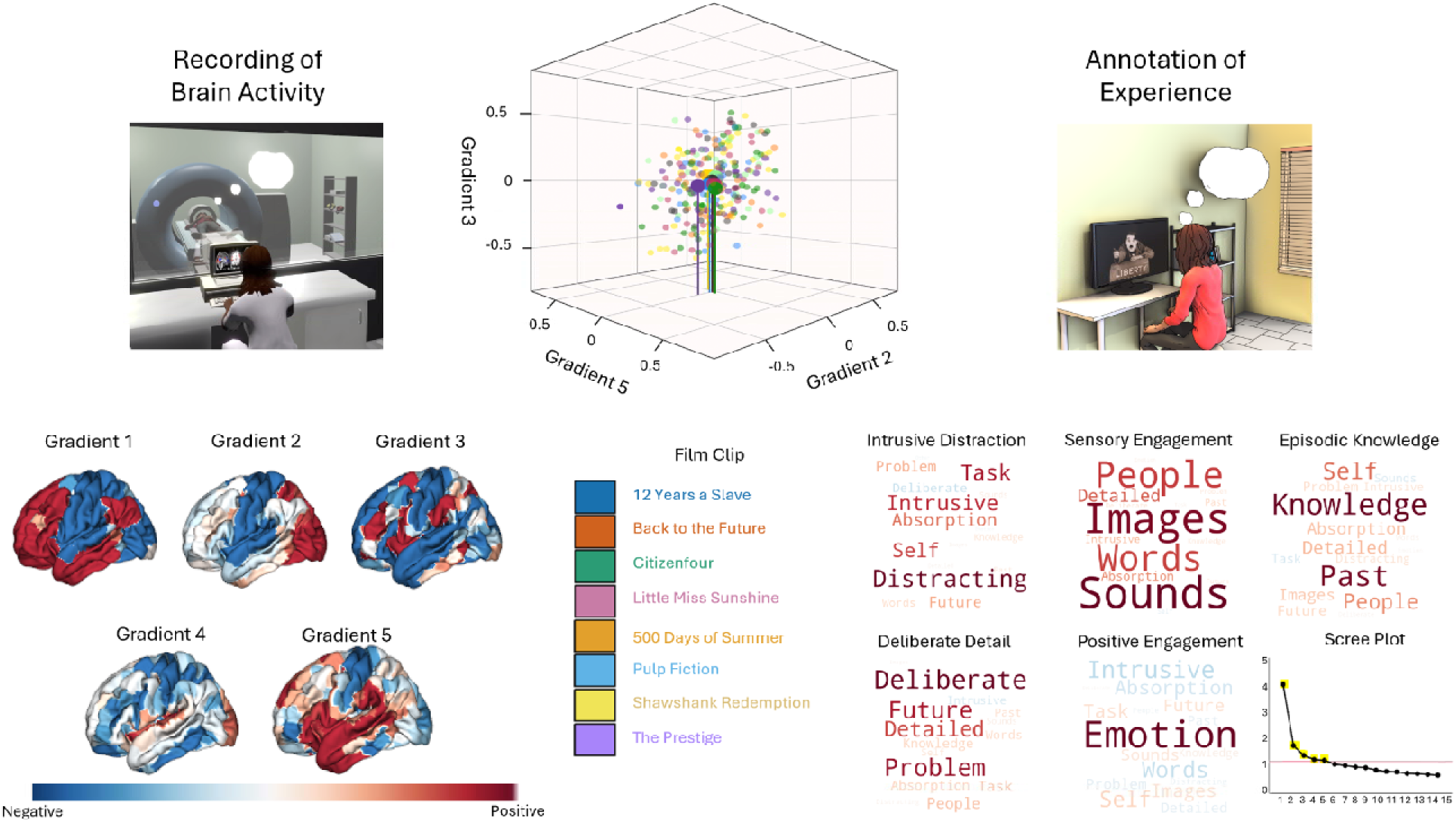
*Note.* Two independent samples contributed uninterrupted recordings of brain activity (fMRI sample, *left*) and annotated experiences (mDES sample, *right*) across eight movies. We used cortical dimensions defined from resting state HCP data (Margulies et al., 2016), which we refer to as gradients (Gradient 1: Primary to Association, Gradient 2: Motor to Visual; Gradient 3: DMN to FPN; Gradient 4: Dorsal Attention Network (DAN) to Visual + Ventral Attention Network (VAN); Gradient 5: Sensorimotor to VAN). Gradient maps were used to calculate gradient coordinates using brain recordings across the films, which are displayed on the bottom left, with Gradient 1-3 maps on top and Gradient 4-5 maps on the bottom (Margulies et al., 2016; McKeown et al., 2025; McKeown et al., 2020; Wallace et al., 2025). The distribution of brain activity across each movie is shown in the top-middle 3D plot (Gradient 5, Gradient 3, Gradient 2), with individual participant scores plotted as smaller points. The colour of the dots corresponds to the film, and the corresponding legend is in the middle. We used Principal Component Analysis (PCA) to decompose the annotated mDES data into patterns of thought. The scree plot from the PCA analysis (bottom middle) provides evidence for a 5-factor component solution, with component number on the x-axis and Eigenvalue on the y-axis. The red line on the y-axis indicates the Eigenvalue threshold of 1. These patterns of thought are shown as word clouds on the bottom right and are *labelled Intrusive Distraction, Sensory Engagement, Episodic Knowledge, Deliberate Detail,* and *Positive Engagement*. In these word clouds, the font size describes their importance (bigger = more important), and the colour describes their polarity (red = positive, blue = negative).

To generate a correspondingly lower-dimensional ‘brain space’, we leveraged the first five dimensions of intrinsic functional connectivity, often referred to as gradients, as derived from a decomposition of diffusion embedding of the Human Connectome Project (HCP) resting-state data (see Methods for further details; see also **Figure 2**; (Margulies et al., 2016; Wallace et al., 2025)). We utilized resting-state gradient maps as gradients obtained during movie-watching have been used to specifically capture functional organization specific to the naturalistic viewing state itself (Samara et al., 2023; Smith et al., 2025). Here, our aim was to characterize thought-brain associations within a coordinate system independent of any single film, facilitating a standardized reference framework across movies and one that remains closely tied to prior gradient-based brain-spaces derived from intrinsic resting-state functional organization (Margulies et al., 2016; Smallwood et al., 2021a). These dimensions describe large-scale connectivity patterns that characterize the topographical organization of brain activity. Data from the group-level averaged BOLD timeseries for each film were projected into the brain space by correlating the whole-brain map at each time point (TR = 1) with each of the first five gradients (see **Figure 1**; (McKeown et al., 2025; McKeown et al., 2023; Wallace et al., 2025)). These correlations then become coordinates that position a momentary state (each TR) within the geometry of the overall brain space (see **Supplementary Figure 1** for illustration of brain space using gradients 1-3 across the duration of each film clip). Gradient 1 describes a functional continuum from primary sensory and motor cortex to heteromodal association cortex. Gradient 2 spans from motor to visual cortex. Gradient 3 ranges from DMN to frontoparietal network (FPN), Gradient 4 spans from dorsal attention network (DAN) to primary visual and ventral attention network (VAN), and Gradient 5 spans from sensorimotor to VAN.

**Figure 2.**
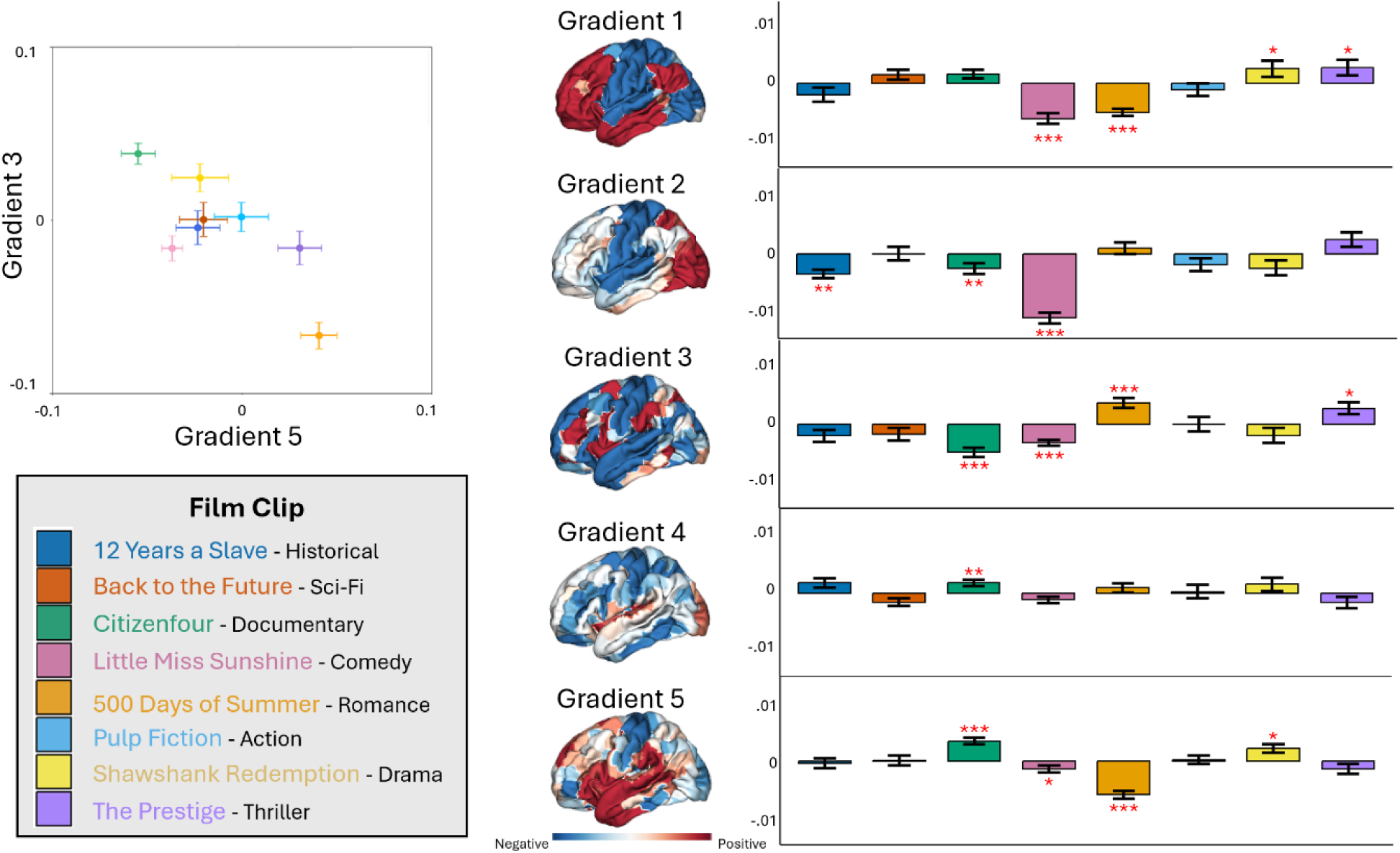
*Note.* Left to right. The heterogeneity of the brain across each of the eight movies are shown in a 2D plot using gradient 5 and gradient 3 (see **Supplementary Figure 2** for additional gradient combination plots). The legend beneath indicates the corresponding movie for each colour in the points in the scatterplot above and the bars in the bar plot to the right. The brain images in the middle represent the activation regions associated with each of the five Gradient maps. The bar plot on the right illustrates the associations for the ANOVA models using Gradients 1-5 as independent variables and each of the movies as dependent variables. The error bars show 95% Confidence Intervals, and the significance asterisk corresponds to the estimated marginal means outcome used to follow up each ANOVA model to test if the gradient effect is significantly different from 0 (* <.05; ** <.01, *** <.001; FDR corrected).

We used our brain spaces, respectively, to determine how heterogeneous brain activity and thought patterns vary across each of the films in our sample. We conducted one-way repeated-measures ANOVAs (Analyses of Variance) with films (8 levels) as the within-subject factor separately for each of the five gradients and each of the five thought components (10 total ANOVAs). We then followed up with Tukey’s HSD pairwise comparisons. This analytical method separates variance attributable to film identity from consistent between-subject differences in group-level average gradient loadings or thought component scores, thus isolating the effect of each film. Lastly, we also computed estimated marginal means (EMMs) for each ANOVA to further test whether each effect was meaningfully different from 0 (see **Figure 2** for the thought space and **Figure 3** for the brain space).

**Figure 3.**
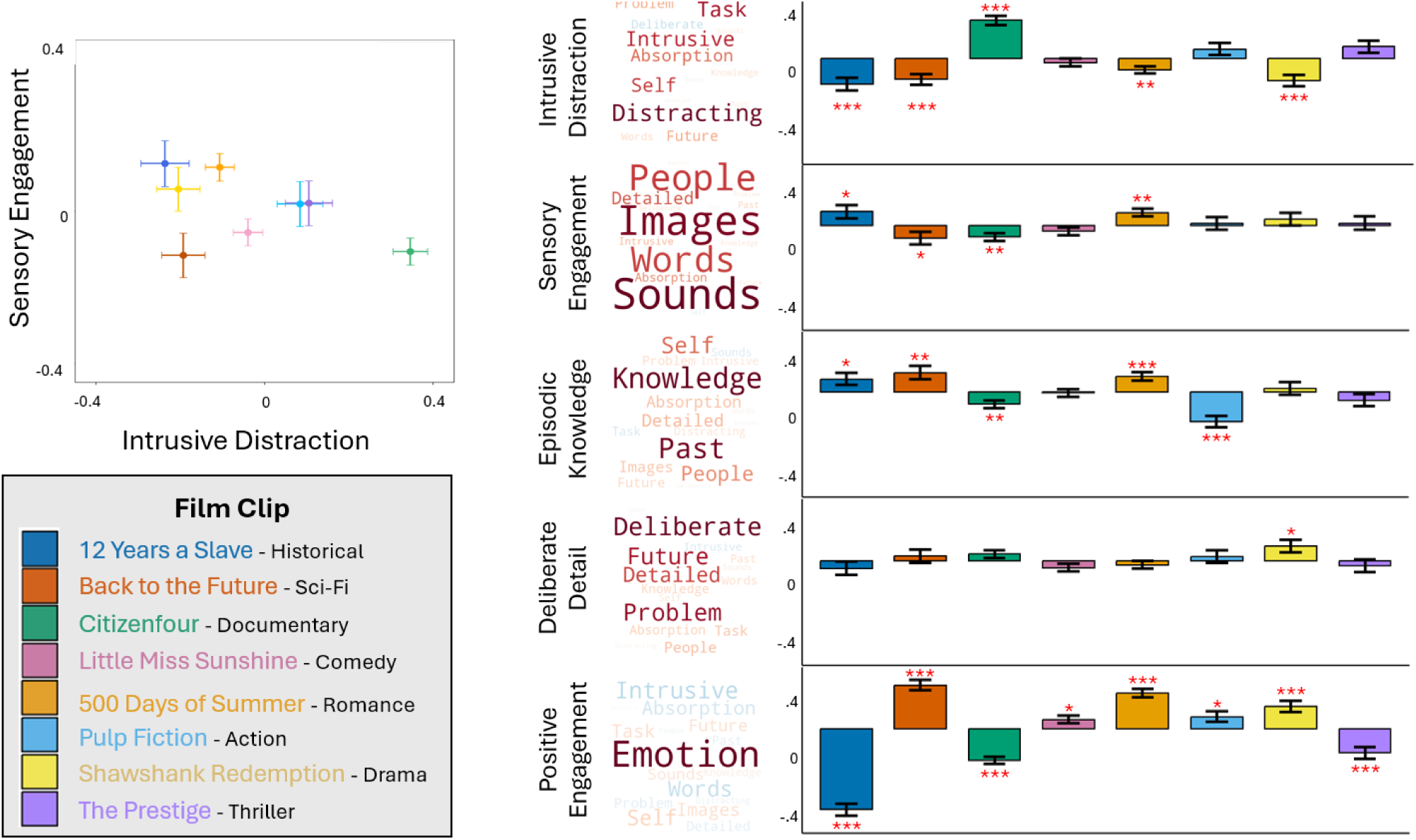
*Note.* Left to right. The heterogeneity of thoughts across each of the eight movies is shown in a 2D plot using *Intrusive Distraction* and *Sensory Engagement* (see **Supplementary Figure 3** for additional thought component combination plots). The legend beneath indicates the corresponding movie for each colour in the points in the scatterplot above and the bars in the bar plot to the right. The bar plot on the right illustrates the associations for the ANOVA models, with each thought component (*Intrusive Distraction, Sensory Engagement, Episodic Knowledge, Deliberate Detail,* and *Positive Engagement*) as an independent variable and each movie as a dependent variable. The error bars show 95% Confidence Intervals, and the significance asterisk corresponds to the estimated marginal means outcome used to follow up each ANOVA model to test if the thought component effect is significantly different from 0 (* <.05; ** <.01, *** <.001; FDR corrected).

Across the brain, all five gradients demonstrated a difference in their associated scores across movies in at least one of the clips (all *p* < .05 FDR corrected; see **Supplementary Table 2** for the ANOVA output; see **Supplementary Table 3** for Tukey’s HSD pairwise comparisons), and EMM analyses further identify which specific films each gradient was reliably present in, marked by the asterisks in **Figure 2** (see also **Supplementary Table 4**). In the thought space, as shown in **Figure 2**, all five thought patterns also varied across movies (all *p* <.05 FDR corrected; see **Supplementary Tables 5-7**), with EMMs again identifying the films for which each thought pattern was reliably expressed (**see Figure 3**).

Together, these results establish that the complex contextual variability across the eight films is evident from both evoked brain activity and subjective experiences across samples. By highlighting various aspects of brain activity and experience, we can visualize how films differ or overlap across these dimensions. (see **Figure 2** and **Figure 3**; see **Supplementary Figures 2-3** for additional combinations of thought components and gradient mappings). Importantly, these findings suggest that films are heterogeneous in both neural and experiential contexts and further underscore the importance of untangling which features of films are specific to the movie itself and which are common across all films.

### 4. Associated Neural-Experiential Features of Movie-Watching

Having established heterogeneity in brain activity and thoughts across movies, we next examined how gradient-based cortical organization related to thought content and whether these associations were robust to variability in movie identity. We conducted two independent sets of linear mixed models predicting each thought pattern from the five gradient maps. In the first set, participant was included as a random intercept; in the second set, both participant and movie identity were included as random intercepts. Comparing the two sets allowed us to ask whether associations between gradient maps and thought content were present when movie identity was, or was not, modelled. We treat this second set as a more conservative test of a stimulus-invariant gradient-thought relationship, since any effect identified there cannot be attributed to any single film in our sample (Yarkoni, 2020).

We thus compared, for each thought pattern, a null model containing only a random intercept for participant against a full model including the five gradient maps as fixed effects and participant as a random intercept (Set 1; Participant-only). We separately compared a null model with random intercepts for both participant and movie identity against a full model that included the five gradient maps as fixed effects and the same two random intercepts (Set 2; Participant + Film). Models within each set were compared using likelihood ratio tests.

Adding the gradients improved fit over the null models for two thought patterns in each set. In the Participant-only models (Set 1), *Intrusive Distraction* showed that including gradient maps significantly improved model fit relative to the null model (χ²(5) = 11.43, p = .044). The same was true for *Sensory Engagement*, where inclusion of gradient maps significantly improved model fit relative to the null model (χ²(5) = 11.11, p = .049). All other models were not significant (*p* > .05; see **Supplementary Table 8** for full model output).

In the Participant + Film models (Set 2), gradients again significantly improved model fit relative to the null for *Sensory Engagement* (χ²(5) = 12.28, p = .031) as well as for *Episodic Knowledge* (χ²(5) = 13.83, p = .017). However, no associations were found for *Intrusive Distraction* or any other thought pattern (*p* > .05; see **Supplementary Table 9** for full model output).

We then examined each gradient’s contribution within the significant Set 1 models (FDR-corrected). *Intrusive Distraction* was negatively predicted by gradient 3 (*F*(1, 3551) = 8.36; *t*(3551) = -2.89, *p* = .019), indicating greater activity towards DMN. *Sensory Engagement* was negatively predicted by gradient 1 (*F*(1, 3543) = 4.12; *t*(3543) = -2.03, *p* = .042), with lower gradient 1 scores indicating greater activity towards sensorimotor cortex. *Sensory Engagement* was also predicted by higher gradient 5 (*F*(1, 3530) = 4.93; *t*(3530) = 2.21, *p* = .026), indicating greater activity in dorsal temporal cortex, auditory cortex, and inferior frontal cortex.

Within the significant models in Set 2 (Participant + Film), we found the gradient 5 effect associated with *Sensory Engagement* was strengthened (*F*(1, 3529) = 9.62; *t*(3529) = 3.10, *p* = .002). Interestingly, despite not appearing in Set 1 (Participant-only), *Episodic Knowledge* was also positively associated with gradient 5 (*F*(1, 3549) = 5.50; *t*(3549) = 2.35, *p* = .019). The full results, including non-significant models, are included in **Supplementary Tables 8-9.** These results are presented in **Figure 4**, with the first set of LMM results in the left-hand column and the second set of LMM results in the right-hand column.

**Figure 4.**
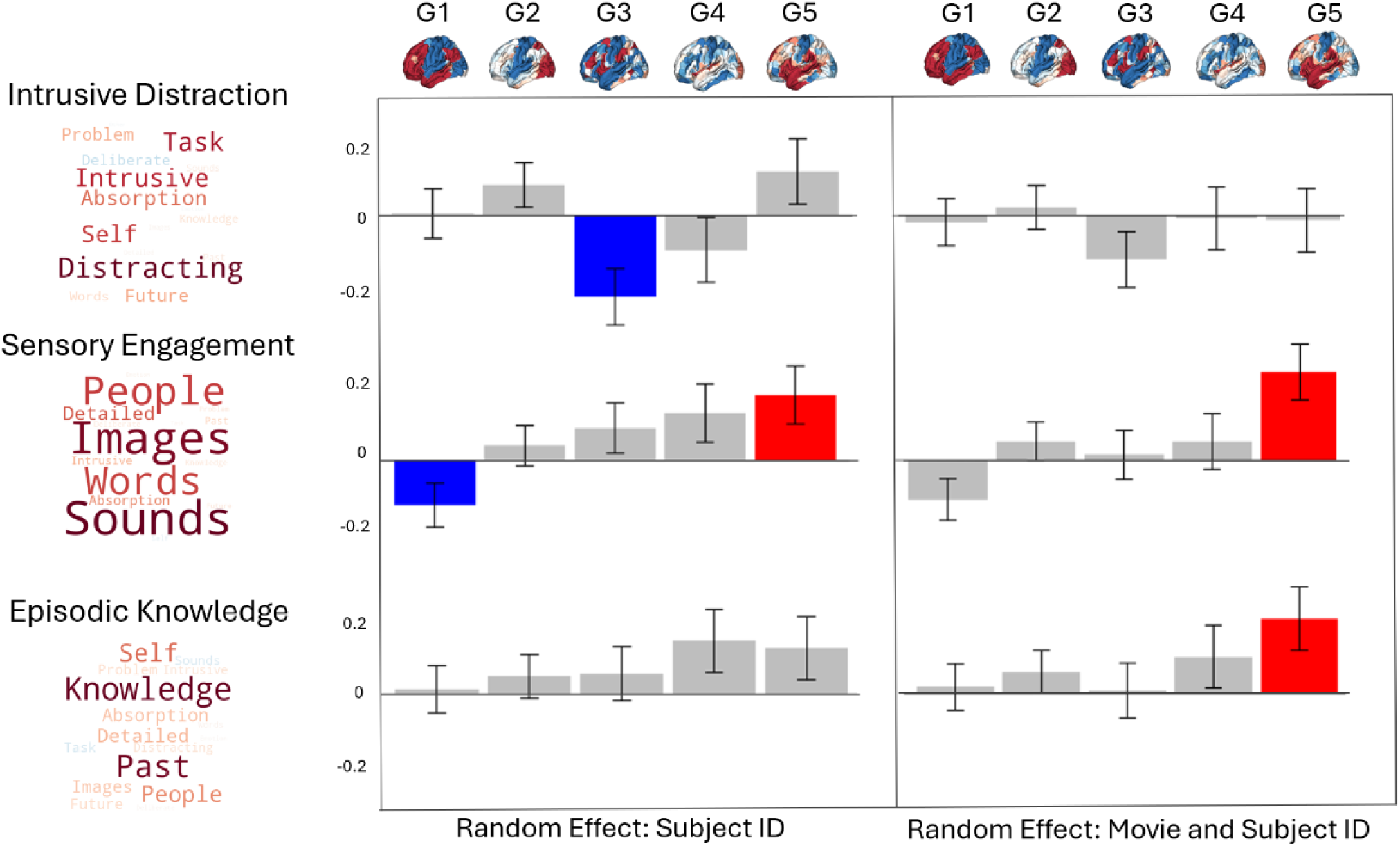
*Note.* Two sets of linear mixed models were computed to predict thought patterns using gradients 1-5 as independent variables. Across the five reported thought patterns in the two sets of models, three reported thought patterns were associated with brain activity (Top: *Episodic Knowledge*, Middle: *Sensory Engagement*, Bottom: *Intrusive Distraction*). The models are identical, except that the parameters on the left correspond to the model that includes participant as a random effect only, and the parameters on the right correspond to the model that includes both participant and movie as random effects. The coloured bars indicate significant parameters and are coloured by effect direction (red = positive, blue = negative). The error bars reflect 95% confidence intervals

### 5. Reconstruction of Movie-Watching Neural Patterns

To visualize the whole-brain states identified by our analysis, we used a dot-product approach that combines functional gradient maps with the LMM beta coefficients from the previous analysis. For each thought pattern in each set of LMMs, excluding intercepts, we extracted beta coefficients for each gradient predictor, whether or not it was significant. Then, for each model, we multiplied each parcel by the sum of its contribution across all gradients, scaled by its associated beta estimate. The result is a single brain map for each thought pattern in each set of LMMs, representing the predicted spatial distribution of brain activity based on the thought-pattern models and their gradient weightings. The maps corresponding to the significant models across each set are displayed in **Figure 5**, as well as their relevant location within the brain space to illustrate the similarities and differences in activation patterns across *Intrusive Distraction* (Set 1; Participant-only), *Sensory Engagement* (Set 2; Participant + Film) and *Episodic Knowledge* (Set 2; Participant + Film). All maps from both sets of LMMs for each thought pattern are shown in **Supplementary Figure 4**.

**Figure 5.**
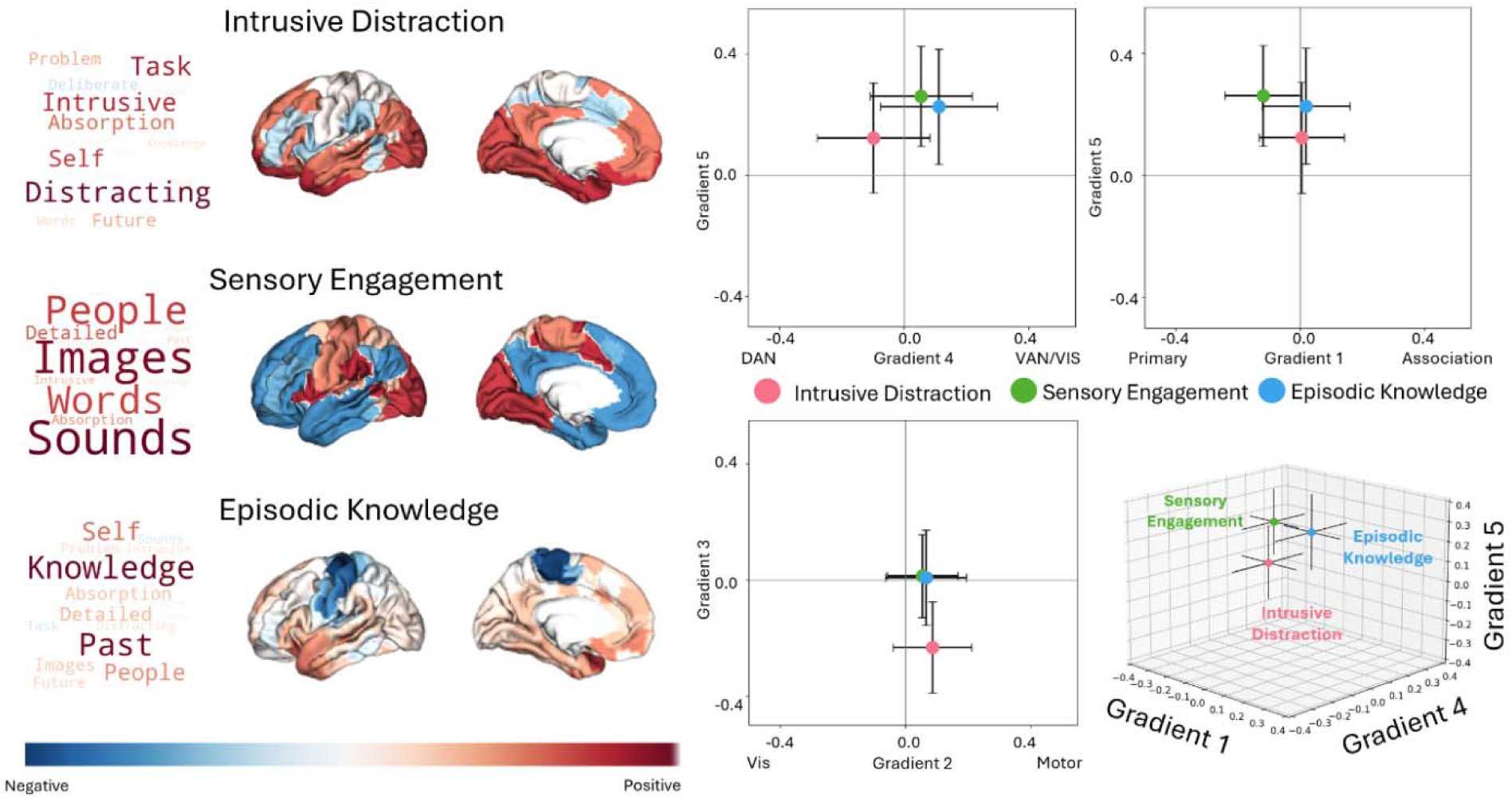
*Note.* Associated brain maps of the thought patterns, *Episodic Knowledge*, *Sensory Engagement*, and *Intrusive Distraction*, were created using the dot-product method using the coefficient estimates from the linear mixed model predicting thought patterns using gradients. The distribution of these associated brain maps is plotted on the right to illustrate the similarities and differences in brain activation patterns across each of the thought patterns. The 3D plot on the bottom right also illustrates the location of the brain across Gradient 1, Gradient 3, and Gradient 4 for the thought patterns *Sensory Engagement* (Green), *Episodic Knowledge* (Blue), and *Intrusive Distraction* (Red).

Next, to further characterize how large-scale brain networks might relate to reported patterns of thought during movie-watching, dot-product values were aggregated and projected onto the Yeo-7 network decomposition derived from resting-state fMRI data (Yeo et al., 2011). The findings are visualized using a bar plot with 95% confidence intervals (see **Figure 6**), which denotes the spatial distribution of the mean values across the seven-network atlas. The cortical surface maps are included for convenience. Our results reveal that *Intrusive Distraction* was associated with positive loadings on the visual network, limbic systems, and the DMN, and negative loadings on the ventral attention network (VAN) and control network. *Sensory Engagement* was associated with positive loadings on the visual, somatomotor, dorsal attention network (DAN), and VAN and negative loadings on limbic, control, and DMN regions. *Episodic Knowledge* was associated with positive loadings on the visual network, VAN, limbic system, and DMN, and negative loadings on the somatomotor network.

**Figure 6.**
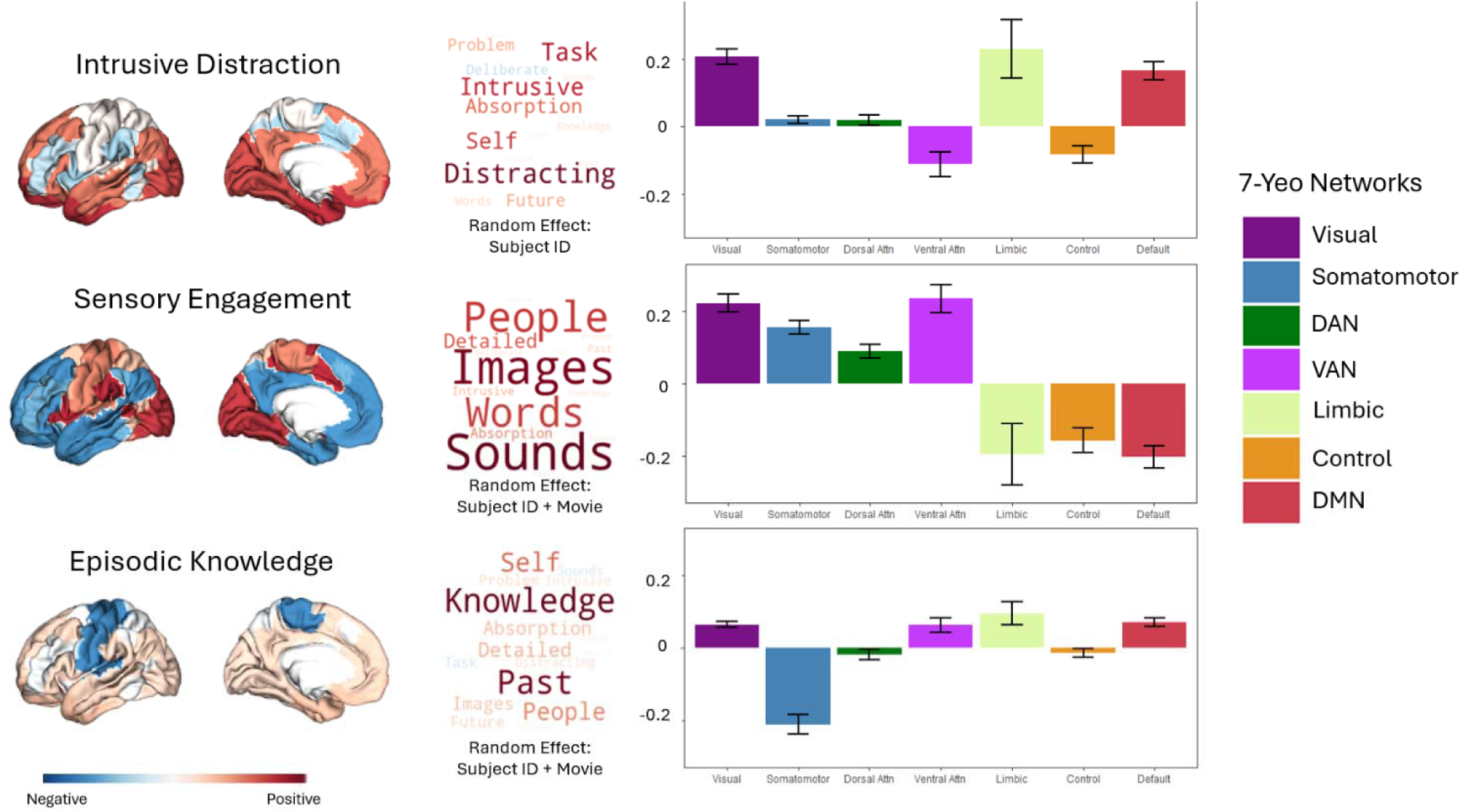
*Note. Left to right.* Mean dot-product values across the Yeo-7 network decomposition are displayed as cortical images, with warmer colours indicating a positive association and cooler colours indicating a negative association. The relevant word clouds for each thought pattern are also included for clarity. The bar plot illustrates the average dot-product between cortical regions and each relevant thought component: *Sensory Engagement* (with movie as a random effect), *Episodic Knowledge* (with movie as a random effect), and *Intrusive Distraction* (without movie as a random effect). The colour bar on the right indicates the corresponding network from the Yeo-7 network.

### 6. Comprehension

Finally, we examined whether participants’ memory for information from the movies could be predicted from their thought patterns. Participants in the laboratory session answered four multiple-choice comprehension questions (four choices) for each movie clip they viewed, based on information from the clip they had just watched. We performed an LMM to predict comprehension scores, using the five thought patterns as independent variables and subject ID as a random effect. The analysis revealed three significant effects. *Intrusive Distraction* negatively predicted comprehension performance (*F*(1, 3735) = 118.03; *t*(3735) = -10.86, *p* < .001), while *Episodic Knowledge* (*F*(1, 3728) = 11.28; *t*(3728) = 3.36, *p* = .001) and *Positive Engagement* (*F*(1, 3660) = 75.10; *t*(3660) = 8.67, *p* <.001) positively predicted comprehension performance. See **Supplementary Table 10** for complete model output.

## 7. Discussion

To our knowledge, this is the first study to combine experience sampling and fMRI across a diverse set of films while modelling films as a random effect, enabling us to test which brain-experiential relationships map within and beyond the movie stimuli sampled. Our study shows that dynamic, multi-dimensional reports of experiential thought during movies can be mapped onto low-dimensional macroscale gradients, revealing how differences in people and films correspond to distinct cognitive-neural states associated with stimulus-driven and context-dependent aspects of films. By measuring experiential reports using multi-dimensional experience sampling (mDES) across eight diverse films, we identified three important features of the movie-watching experience that mapped to the brain: (i) perceptually-coupled engagement with visual and auditory features of the movie (*Sensory Engagement*), (ii) situations where prior knowledge is brought to bear to aid comprehension of the movie (*Episodic Knowledge*) and (iii) patterns of distracted thought that were linked to poor comprehension of the film (*Intrusive Distraction*). By leveraging continuous cortical gradients from the Human Connectome Project (HCP), we mapped each of these states onto whole-brain patterns of functional organization (Margulies et al., 2016; Van Essen et al., 2012). Together, our analyses indicate that, during naturalistic viewing, *Sensory Engagement* reflects a robust, perceptually coupled state anchored in sensory cortex (visual and auditory) and attention systems. Higher-order states, such as *Episodic Knowledge* and *Intrusive Distraction*, despite their opposing relationship with comprehension, were both associated with increased activation in limbic and default-mode regions (Kalisch et al., 2006; McKeown et al., 2025; Ritchey, 2026). A unique advantage of our analysis is that using gradient coordinates allows us to situate whole-brain activity along established, continuous dimensions of cortical organization. Importantly, this approach allows us to parse the functional heterogeneity of the DMN observed across these two contrasting states by looking at their locations in brain space: While *Intrusive Distraction* and *Episodic Knowledge* both engage whole-brain states with increased DMN activity, they are distinguishable based on their unique relationships with other networks, such as the negative association of gradient 3 and the positive association of gradients 4 and 5. Together, these findings highlight how thought-brain spaces can capture a hierarchical organization of distinct, even opposing, cognitive states, such as sensory and memory-associated systems, and how the same functional networks can facilitate this organization despite their different characterizations in reported experience.

A critical contribution of this work is the ability to distinguish between intrinsic domain-general cognitive mechanisms and stimulus-specific effects – a challenge recently highlighted in naturalistic neuroscience research (Grall & Finn, 2022; Leipold et al., 2025). To address this challenge, our approach drew on eight films across eight genres to examine diverse experiences and elicit a high degree of variability in movie-watching stimuli. Then, we employed two analytical approaches to partition the variance related to individual differences and movie stimuli: (1) one set, with participants as a random effect, which identifies associations between experiential aspects across the eight specific films in our sample, and (2) a second set, with participants and movie as random effects. In this way, our approach aimed to understand how thoughts differ across contexts and what variance might be attributed to neural mechanisms, regardless of the person or the film. By integrating stimulus variables, such as the movie as a random intercept effects, similar to how we control for variability among participants, we identify which brain-experiential mappings are most robust to contextual variability and therefore likely represent a general rule about how thoughts are instantiated in the brain during movie watching. In contrast, findings from the first set of models, with only participant variability controlled for, likely reflect thought-brain mappings influenced by specific experiential features of the films in our sample and demonstrate how contextually narrative features drive variability in experiential states. Collectively, by interpreting the outcome of the effects in both models relative to experience, we can better understand the robustness of our effects across neural activation patterns in films.

*Sensory Engagement* was the most stable feature of movie-watching, identified in both sets of models, and suggests that the link between sensory-related thoughts and the sensory cortex is the most robust feature of the movie-watching experience and is unaffected by variations in narrative context. Additionally, the neural association with *Episodic Knowledge*, a state that supports better comprehension performance, emerged only after controlling for movie-level variance. These results suggest that although the neural mechanism linking thoughts about the past, knowledge, and narrative integration is consistent across films, it is not detectable until film-specific variability is accounted for. In other words, certain elements of films may obscure this state, but once we account for the noise across movies, the underlying stimulus-invariant mechanism emerges. Conversely, *Intrusive Distraction* was associated only with the first set of models, suggesting that this brain–thought association is contingent on film-level variance: it is detectable when that variance is left unmodelled but not once it is accounted for. Our results suggest that *Intrusive Distraction* likely describes unsuccessful narrative anchoring (Smallwood, 2011; Smallwood et al., 2008; Steindorf et al., 2023). Unlike stimulus-coupling mechanisms, this context-dependent state reflects disengagement from the stimulus, associated with reduced comprehension performance, and is likely driven by unique features of films rather than by a general mechanism of film watching. These findings suggest that the DMN plays a flexible role during movie-watching, likely connected to control systems. Studies on reading demonstrate that narrative comprehension relies on perceptual coupling within the DMN, whereas perceptual decoupling leads to mind-wandering and reduced comprehension (Medea et al., 2018; Zhang et al., 2022). However, our results show that DMN can support both improved and impaired memory performance depending on perceptually coupled states that differ in engagement with attention and control systems (Smallwood et al., 2021b; Wallace et al., 2025).

Our analysis has several practical implications for understanding brain activity during films, since differences across films obscure or alter the meaning of any psychological links with brain activity. Movie-watching paradigms provide a rich, multimodal, and dynamic framework for investigating how cognition unfolds in real-world applications while maximizing experimental control that is not possible with fully unconstrained paradigms (Nastase et al., 2019; Zhang et al., 2021). Furthermore, by leveraging movie-watching neuroimaging in our work, we can better understand how contextual narrative features affect brain activity across lower- and higher-order cortical regions through mDES. Based on our analysis, many brain-behaviour associations may vary in strength across films. There may also be more complex dependencies that reflect how different groups of people respond to different films, as existing neuroimaging research has highlighted the utility of movie-watching paradigms for predicting cognitive, emotional, and even clinical features in individuals (notably outperforming rest) (Eickhoff et al., 2020; Finn & Bandettini, 2021; Gal et al., 2022; Grall & Finn, 2022; Vanderwal et al., 2015). Consistent with these applications, associations between personality traits and brain activity derived from task data from the HCP reflect task-specific changes rather than a consistent, chronic influence across all situations (Hardikar et al., 2024). The most obvious solution to the heterogeneity of brain activity across films is for studies to use a sufficiently wide range of movie stimuli, allowing superficial differences to be removed via random effects. An alternative would be to develop theoretical models of movie-watching that account for film features (e.g., genre) using techniques such as mDES, and to use these models to guide stimulus selection for more expensive, resource-intensive scanning studies. This approach would allow large corpora of film stimuli to be curated based on their experiential features using mDES (a relatively inexpensive, scalable, and efficient technique), while also measuring traits of interest. This efficient approach identifies movies that best discriminate among traits or variables of interest, thereby enabling more precise application in resource-intensive brain imaging experiments or clinical interventions. Without such methodological rigour, researchers should remain explicitly cautious when interpreting mappings between neural activity and the psychological features of specific films.

Our study advances our understanding of the movie-watching experience and how psychological function maps onto different neural systems across naturalistic contexts. Our study suggests that, at its core, the movie-watching experience is a sensory state that anchors the individual in a perceptually coupled mode, amenable to analysis of the auditory-visual features that constitute a film. This was the most consistent experiential-brain mapping identified in our analysis, suggesting that it is likely a core element of almost all movie-watching experiences. However, successful understanding and comprehension of the film requires more than just sensory coupling. *Episodic Knowledge* was associated with better comprehension and rebalancing of neural activity towards memory systems and away from sensory-motor regions, but only once film-level variance was modelled. More broadly, our findings suggest that in naturalistic designs, variability between films is not simply noise to be averaged away but part of what determines which links between brain and experience we can observe.

## 8. Methods

### Participant pool

The sample consisted of 226 participants (185 women (81.5%), 34 men (15%), six non-binary (2.6%), one other gender identity (<1%), one prefer not to say (<1%); age: *M* = 19.03, *SD±* 1.30, (17-30)) who participated in the in-person laboratory study to watch the short movie clips, responded to mDES probes about their thoughts, and completed a brief comprehension quiz on information in the film clips. All participants spoke English, and the majority of the sample primarily resided in Canada (<97%; China (n = 3), India (n = 2), Nigeria (n = 1), USA (n = 2). This study was granted ethics clearance by the Queen’s University General Research Ethics Board (#6036804). Participants were recruited between March 2023 and April 2024 through the Queen’s University Psychology Participant Pool. Participants read and provided informed written consent via Qualtrics before participating in the research study. Participants were rewarded with one-course credit or $10.00 CAD for their participation and were provided with a verbal and written debrief form upon completion of the study.

### Participant pool – brain data sample

The sample included in the neuroimaging data consisted of 73 participants (35 Female, 38 male; *M±SD_age_*= 27.63 ±10.82 years, range = 18-58 years). See Aliko and colleagues for further description of the sample (Aliko et al., 2020).

### Procedure

Participants were recruited from Queen’s University and attended a single, 1-hour testing session in the laboratory. After providing written and informed consent, participants voluntarily provided non-identifying demographic information, such as their age, gender, sex, language, and country of residence. Participants were registered and assigned to a computer testing booth, where they were seated at a desk and provided with a pair of headphones to listen to and attend to the movie clips on the computer screen. Participants were randomly assigned to a subset of the clips to watch in a randomized presentation. During the presentation of each movie clip, participants were briefly interrupted five times, approximately two minutes apart (jittered), to answer mDES probes about the content of their thoughts. Participants were assigned to counterbalanced, unique probe orders to ensure that probes were delivered in a pseudo-random manner across the presentation of each clip. Probe orders were not repeated per participant. The first probe was delivered at least 60 seconds after the clip began. After each clip concluded, and all the mDES probes were responded to, participants briefly answered four questions about the information in the movie clip they had just watched before the following clip began. Once all the clips had been watched and the questionnaires had been responded to, participants were debriefed and awarded compensation of their choice.

### Multi-dimensional experience sampling (mDES)

Participants received a total of 15 mDES probes across the three clips, with five probes for each, and all responses were made with respect to their thoughts immediately before the probe interrupted their viewing. No probes were administered within the first minute of the clip — the first possible probe was administered at the 70-second mark, allowing participants to situate themselves within the context of the movie clip. Each of the 16 mDES questions appeared in a randomized order, and participants were asked to use the directional arrow keys to move a slider across the screen to indicate, on a scale of 1 (not at all) to 10 (completely), how much that particular feature characterized their thoughts by pressing the enter key. The specific items used in this experiment are presented in Table 1.

Probe orders were created using Python to ensure sufficient spacing between probes and prevent probe durations from repeating by using parameters such as the number of clips in the battery, the duration of the clip for probe delivery, the frequency of probe delivery at the individual and group level, the minimum time between probes, and the number of participants at each probing interval. This enables us to create a counter-balanced, randomly assigned probe condition for each participant, ensuring that each participant is assigned to a different probe order for each clip they watch. Across the sample, this allows us to determine how many participants are needed to fulfill the specified parameters for the probe orders to equally balance the number of mDES observations across the entire clip. This method allows us to sample experience as frequently as possible without interrupting the natural viewing process that would otherwise interrupt the movie-watching experience. Furthermore, this approach ensures that participants themselves are not probed too frequently about their experience, while maximizing the possible observations we can acquire from unique subset combinations of participants, virtually minimizing participant biases. The corresponding analysis code is available at Mendeley Data.

### Movie Clip Stimuli

Movie clips were presented in 9-to-11-minute scenes from *Citizenfour*, *Little Miss Sunshine*, *500 Days of Summer, Back to the Future, The Prestige, Pulp Fiction, 12 Years a Slave, and The Shawshank Redemption.* We selected these films from the Naturalistic Neuroimaging Database (NNDb; (Aliko et al., 2020)), where participants viewed them in an fMRI scanner. Our laboratory sample of participants was informed that they would watch a subset of these movie clips (3 or 5 of the clips) from different genres of film presented in a random order. Instructions were provided to participant on screen to attend to the stimuli, respond to mDES probes, and answer the comprehension questions.

### Comprehension

Following each clip assigned to the participant, a short 4-question multiple-choice comprehension quiz was immediately presented. Participants were instructed not to take any notes or prepare for the comprehension questions. These questions were collaboratively created to test knowledge of the movie content or events in the clip, particularly, though without requiring study or note-taking. Additionally, questions were designed to elicit information about events that occurred during the beginning, middle, and end of the clip. An example of a comprehension question for the movie clip *Back to the Future* was “How much is the cup of coffee advertised for in the diner?” in which participants could choose a) 5 cents, b) $1.00, c) $3.50, or d) $4.50. Participants answered the question by reading the multiple-choice options and indicated their response using the corresponding letter on the keyboard. A table of all the questions, with the multiple-choice options and corresponding answers, can be found in **Supplementary Table 11**.

### Experiential Analysis

Participants in our laboratory responded to multi-dimensional experience sampling (mDES) probes about features of their experience while watching movie clips. To create the thought patterns used in our analyses, we derive all the experiential reports from participants across all movies to perform a data reduction analysis using Principal Components Analysis. We retained components based on the scree plot and the eigenvalue threshold of 1.

### Brain Analysis

Our analyses used brain data acquired and shared by the NNDb, an open-access database of pre-processed MNI 2 mm fMRI data (TR = 1) of participants who watched one of 10 full-length movies (Aliko et al., 2020). We utilized MNI 2 mm fMRI data from the NNDb open database from participants who watched Citizenfour (n=18), Little Miss Sunshine (n=6), 500 Days of Summer (n=20), Back to the Future (n = 6), The Prestige (n = 6), Pulp Fiction (n = 6), 12 Years a Slave (n = 6), and The Shawshank Redemption (n = 6). The specific pre-processing steps applied to the brain data and specific details of the sample are described elsewhere (Aliko et al., 2020); briefly, the fMRI data were slice-timing and motion corrected, non-linearly normalized to the MNI template, spatially smoothed using a 6 mm FWHM Gaussian kernel, detrended against motion, white matter, and cerebrospinal fluid signal regressors, and cleaned of ICA-identified artifacts.

Our analysis derived the pre-processed data from Aliko and colleagues (Aliko et al., 2020). We first extracted the brain activity for each individual from the 8- to 10-minute section that we sampled for using mDES to accurately map our mDES timeseries from our laboratory sample onto the corresponding brain data from the NNDb sample. This created a time series of TR scores for every second in the movie clips we used in our experiment by taking the group-level average across TRs. From here, we projected these group-level TR time series onto the first five dimensions of the gradient maps, which were produced from a non-linear eigenvector decomposition of resting-state fMRI connectivity data (Margulies et al., 2016) from the Human Connectome Project (HCP) dataset (Van Essen et al., 2012). Each probe response was matched to the corresponding gradient correlate at the TR coincident with probe onset and interpolated between probes to understand the neural composition of brain regions derived from the fMRI data obtained from the NNDb sample for each movie clip.

### Heterogeneity Analysis

Two heterogeneity analyses were computed to understand the diversity in thought and brain expression across the eight films. Analysis of variance (ANOVA) models were computed to assess whether group-averaged scores across the five gradients varied across the eight films. In other words, this model compared the scattering of all the TRs expressed in five gradient coordinates from the mean timeseries and then tested whether the point clouds differed across the different movies. Significant ANOVA models were followed up with pairwise comparisons using Tukey’s HSD. In addition, estimated marginal means (EMMs) models were also conducted to assess if the average gradient score for each film was significantly different from 0. Another set of ANOVA models was computed to assess whether group-averaged scores across the five thought components (*Intrusive Distraction*, *Sensory Engagement*, *Episodic Knowledge*, *Deliberate Detail*, and *Positive Engagement*) varied across the eight films. Similarly, significantly ANOVA models were followed up with pairwise comparisons using Tukey’s HSD and EMMs to assess if scores significantly differed from 0. FDR was used to account for multiple comparisons.

### Comprehension Analysis

Comprehension was analyzed using a linear mixed model (LMM) to predict comprehension scores across the films, with the five thought patterns as independent variables and subject ID as a random effect. Bonferroni was used to control family-wise error (FWE).

### Thought-Brain Analysis

To assess if the gradient scores were associated with reported thought patterns, we conducted a series of LMMs to predict thought components using Gradients 1-5 as predictors. We computed two sets of models: one that included only subject ID as a random effect and one that included subject ID and movie as random effects. FDR was used to control for multiple comparisons. The first set of models, with only subject ID as a random effect, suggests that any significant findings may be associated with the unique features of the eight films we used in our experiment. The second set of models, with subject ID and movie as random effects, may suggest that any significant findings are associated with movie-watching regardless of the film shown. This modelling structure considers the perspective of Yarkoni regarding the generalizability crisis, which suggests that stimuli in experiments should be treated similarly to intersubject variability in order to better understand their contributions in analytical interpretation (Yarkoni, 2020).

## Acknowledgements

RSW acknowledges support from the Natural Sciences and Engineering Research Council of Canada (NSERC), through a Canada Graduate Scholarship-Doctoral (CGS D – 601452 - 2025). DSM acknowledges support from the European Research Council (ERC) under the European Union’s Horizon 2020 research and innovation programme (grant agreement No. 866533-CORTIGRAD). BCB acknowledges support from the Canadian Institutes of Health Research, CIHR (FDN-154298, PJT-174995, PJT-206196, CIHR PJT-203761, CIHR PJT-191853), SickKids Foundation (NI17-039), Natural Sciences and Engineering Research Council (NSERC RGPIN-2025-05932), Azrieli Center for Autism Research of the Montreal Neurological Institute (ACAR), BrainCanada, FRQ-S, the Helmholtz International BigBrain Analytics and Learning Laboratory (Hiball), Healthy Brains and Healthy Lives (HBHL), Centre for Aging + Brain Health Innovation (CABHI), the Canada Research Chairs Program (CRC), and the Centre of Excellence in Epilepsy at the Neuro (CEEN). JS acknowledges support from Natural Sciences and Engineering Research Council of Canada (#RGPIN 2023-03496).

