## Supplementary Materials for "Brain-thought states during movie-watching reflect shared sensory anchors and film- and person- specific processes in association cortex"

**Supplementary Table 1 – Varimax Rotated Component Matrix**

| mDES Item | Component 1  *Intrusive Distraction* | Component 2  *Sensory Engagement* | Component 3  *Episodic Knowledge* | Component 4  *Deliberate Detail* | Component 5  *Positive Engagement* |
| --- | --- | --- | --- | --- | --- |
| Task | .654 | .052 | -.135 | .193 | .150 |
| Self | .549 | -.089 | .457 | .089 | .205 |
| People | -.025 | .520 | .383 | .210 | -.045 |
| Knowledge | .111 | .100 | .811 | .154 | .061 |
| Absorption | .439 | .357 | .235 | .161 | -.169 |
| Words | .159 | .509 | .044 | .130 | -.216 |
| Past | .074 | .170 | .783 | .118 | -.079 |
| Deliberate | -.190 | .100 | .027 | .705 | .015 |
| Sounds | .071 | .745 | -.118 | .060 | .115 |
| Future | .305 | .094 | .168 | .585 | .137 |
| Problem | .292 | .093 | .147 | .619 | -.097 |
| Distracting | .787 | .004 | .117 | .034 | -.066 |
| Images | .031 | .704 | .219 | .026 | .146 |
| Emotion | -.023 | .050 | .023 | .006 | .906 |
| Intrusive | .604 | .256 | .114 | -.115 | -.243 |
| Detailed | .032 | .461 | .272 | .474 | -.087 |

*Note.* Varimax Rotated Component Matrix for 5 Component Decomposition using *mDES* reports to decompose data into patterns of thought components.

**Supplementary Table 2 – ANOVA (Gradients)**

| Gradient 1 | df | Sum Sq | Mean Sq | *F* value | *p* value |
| --- | --- | --- | --- | --- | --- |
| Task_name | 7 | 5.72 | 0.8168 | 14.82 | <.001 |
| Residuals | 3733 | 205.75 | 0.0551 |  |  |
| Gradient 2 |  |  |  |  |  |
| Task_name | 7 | 8.93 | 1.276 | 19.05 | <.001 |
| Residuals | 3733 | 249.98 | 0.067 |  |  |
| Gradient 3 |  |  |  |  |  |
| Task_name | 7 | 4.42 | 0.6321 | 12.39 | <.001 |
| Residuals | 3733 | 190.18 | 0.051 |  |  |
| Gradient 4 |  |  |  |  |  |
| Task_name | 7 | 0.87 | 0.12441 | 3.389 | .001 |
| Residuals | 3733 | 137.06 | 0.03672 |  |  |
| Gradient 5 |  |  |  |  |  |
| Task_name | 7 | 4.43 | 0.6332 | 20.12 | <.001 |
| Residuals | 3733 | 117.46 | 0.0315 |  |  |

*Note.* ANOVA Model Outcome using Task_name (movie variable) to assess if a difference exists in Gradient scores across the movies. Significant results for the main effect of Task_name suggests there is at least one difference in Gradient scores in one of the films.

**Supplementary Table 3 – Tukey’s HSD Pairwise Comparisons (Gradients)**

| **Gradient 1** | diff | lwr | upr | *p*-val |
| --- | --- | --- | --- | --- |
| back to the future-12 years | 0.039 | -0.017 | 0.095 | .402 |
| citizenfour - 12 years | 0.041 | -0.007 | 0.089 | .163 |
| little miss sunshine - 12 years | -0.047 | -0.094 | 0.001 | .064 |
| summer - 12 years | -0.034 | -0.082 | 0.014 | .366 |
| pulp fiction - 12 years | 0.010 | -0.047 | 0.066 | .999 |
| shawshank - 12 years | 0.051 | -0.005 | 0.107 | .103 |
| prestige - 12 years | 0.053 | -0.003 | 0.109 | .083 |
| citizenfour - backtothefuture | 0.001 | -0.046 | 0.049 | .999 |
| little miss sunshine - backtothefuture | -0.086 | -0.134 | -0.038 | <.001 |
| summer - backtothefuture | -0.074 | -0.122 | -0.026 | <.001 |
| pulp fiction - backtothefuture | -0.030 | -0.086 | 0.027 | .752 |
| shawshank - backtothefuture | 0.012 | -0.044 | 0.068 | .998 |
| prestige - backtothefuture | 0.014 | -0.043 | 0.070 | .996 |
| little miss sunshine - citizenfour | -0.087 | -0.125 | -0.050 | <.001 |
| summer - citizenfour | -0.075 | -0.113 | -0.037 | <.001 |
| pulp fiction - citizenfour | -0.031 | -0.079 | 0.017 | .505 |
| shawshank - citizenfour | 0.011 | -0.037 | 0.058 | .998 |
| prestige - citizenfour | 0.012 | -0.036 | 0.060 | .995 |
| summer - little miss sunshine | 0.012 | -0.026 | 0.050 | .978 |
| pulp fiction - little miss sunshine | 0.056 | 0.008 | 0.104 | .009 |
| shawshank - little miss sunshine | 0.098 | 0.050 | 0.146 | <.001 |
| prestige - little miss sunshine | 0.099 | 0.052 | 0.147 | <.001 |
| pulp fiction - summer | 0.044 | -0.004 | 0.092 | .098 |
| shawshank - summer | 0.086 | 0.038 | 0.134 | <.001 |
| prestige - summer | 0.087 | 0.039 | 0.135 | <.001 |
| shawshank - pulp fiction | 0.042 | -0.015 | 0.098 | .325 |
| pretige - pulp fiction | 0.043 | -0.013 | 0.099 | .279 |
| prestige - shawshank | 0.002 | -0.055 | 0.058 | .999 |
| **Gradient 2** | diff | lwr | upr | *p*-val |
| back to the future-12 years | 0.039 | -0.023 | 0.101 | .543 |
| citizenfour - 12 years | 0.010 | -0.043 | 0.063 | .999 |
| little miss sunshine - 12 years | -0.086 | -0.138 | -0.033 | <.001 |
| summer - 12 years | 0.050 | -0.003 | 0.102 | .085 |
| pulp fiction - 12 years | 0.018 | -0.044 | 0.080 | .989 |
| shawshank - 12 years | 0.011 | -0.051 | 0.073 | .999 |
| prestige - 12 years | 0.066 | 0.004 | 0.128 | .026 |
| citizenfour - backtothefuture | -0.029 | -0.082 | 0.024 | .711 |
| little miss sunshine - backtothefuture | -0.125 | -0.177 | -0.072 | <.001 |
| summer - backtothefuture | 0.010 | -0.042 | 0.063 | .999 |
| pulp fiction - backtothefuture | -0.021 | -0.083 | 0.040 | .966 |
| shawshank - backtothefuture | -0.028 | -0.090 | 0.034 | .870 |
| prestige - backtothefuture | 0.027 | -0.035 | 0.089 | .884 |
| little miss sunshine - citizenfour | -0.096 | -0.137 | -0.054 | <.001 |
| summer - citizenfour | 0.039 | -0.002 | 0.081 | .079 |
| pulp fiction - citizenfour | 0.007 | -0.045 | 0.060 | .999 |
| shawshank - citizenfour | 0.001 | -0.052 | 0.054 | .999 |
| prestige - citizenfour | 0.056 | 0.004 | 0.109 | .027 |
| summer - little miss sunshine | 0.135 | 0.094 | 0.177 | <.001 |
| pulp fiction - little miss sunshine | 0.103 | 0.050 | 0.156 | <.001 |
| shawshank - little miss sunshine | 0.097 | 0.044 | 0.149 | <.001 |
| prestige - little miss sunshine | 0.152 | 0.099 | 0.205 | <.001 |
| pulp fiction - summer | -0.032 | -0.085 | 0.021 | .596 |
| shawshank - summer | -0.039 | -0.091 | 0.014 | .345 |
| prestige - summer | 0.017 | -0.036 | 0.070 | .979 |
| shawshank - pulp fiction | -0.007 | -0.068 | 0.055 | .999 |
| pretige - pulp fiction | 0.049 | -0.013 | 0.111 | .246 |
| prestige - shawshank | 0.055 | -0.007 | 0.117 | .119 |
| **Gradient 3** | diff | lwr | upr | *p*-val |
| back to the future-12 years | 0.003 | -0.051 | 0.057 | .999 |
| citizenfour - 12 years | -0.032 | -0.078 | 0.014 | .430 |
| little miss sunshine - 12 years | -0.014 | -0.060 | 0.032 | .986 |
| summer - 12 years | 0.064 | 0.018 | 0.110 | .001 |
| pulp fiction - 12 years | 0.023 | -0.031 | 0.077 | .903 |
| shawshank - 12 years | 0.001 | -0.053 | 0.055 | .999 |
| prestige - 12 years | 0.054 | 0.000 | 0.108 | .053 |
| citizenfour - backtothefuture | -0.034 | -0.080 | 0.012 | .311 |
| little miss sunshine - backtothefuture | -0.017 | -0.063 | 0.029 | .959 |
| summer - backtothefuture | 0.061 | 0.015 | 0.107 | .002 |
| pulp fiction - backtothefuture | 0.020 | -0.034 | 0.074 | .951 |
| shawshank - backtothefuture | -0.002 | -0.056 | 0.052 | .999 |
| prestige - backtothefuture | 0.051 | -0.003 | 0.105 | .084 |
| little miss sunshine - citizenfour | 0.018 | -0.018 | 0.054 | .808 |
| summer - citizenfour | 0.095 | 0.059 | 0.132 | <.001 |
| pulp fiction - citizenfour | 0.054 | 0.008 | 0.100 | .008 |
| shawshank - citizenfour | 0.033 | -0.013 | 0.079 | .382 |
| prestige - citizenfour | 0.085 | 0.039 | 0.131 | <.001 |
| summer - little miss sunshine | 0.077 | 0.041 | 0.114 | <.001 |
| pulp fiction - little miss sunshine | 0.037 | -0.009 | 0.083 | .236 |
| shawshank - little miss sunshine | 0.015 | -0.031 | 0.061 | .979 |
| prestige - little miss sunshine | 0.067 | 0.021 | 0.113 | <.001 |
| pulp fiction - summer | -0.041 | -0.087 | 0.005 | .125 |
| shawshank - summer | -0.063 | -0.109 | -0.017 | .001 |
| prestige - summer | -0.010 | -0.056 | 0.036 | .998 |
| shawshank - pulp fiction | -0.022 | -0.076 | 0.032 | .924 |
| pretige - pulp fiction | 0.031 | -0.023 | 0.085 | .673 |
| prestige - shawshank | 0.053 | -0.002 | 0.107 | .064 |
| **Gradient 4** | diff | lwr | upr | *p*-val |
| back to the future-12 years | -0.038 | -0.084 | 0.008 | .199 |
| citizenfour - 12 years | 0.000 | -0.039 | 0.039 | .999 |
| little miss sunshine - 12 years | -0.033 | -0.072 | 0.006 | .174 |
| summer - 12 years | -0.010 | -0.049 | 0.029 | .995 |
| pulp fiction - 12 years | -0.017 | -0.063 | 0.029 | .954 |
| shawshank - 12 years | -0.002 | -0.048 | 0.044 | .999 |
| prestige - 12 years | -0.038 | -0.084 | 0.007 | .177 |
| citizenfour - backtothefuture | 0.038 | -0.001 | 0.077 | .066 |
| little miss sunshine - backtothefuture | 0.005 | -0.034 | 0.044 | .999 |
| summer - backtothefuture | 0.028 | -0.011 | 0.067 | .369 |
| pulp fiction - backtothefuture | 0.021 | -0.025 | 0.067 | .866 |
| shawshank - backtothefuture | 0.035 | -0.011 | 0.081 | .274 |
| prestige - backtothefuture | -0.001 | -0.047 | 0.045 | .999 |
| little miss sunshine - citizenfour | -0.033 | -0.064 | -0.002 | .026 |
| summer - citizenfour | -0.010 | -0.041 | 0.021 | .979 |
| pulp fiction - citizenfour | -0.017 | -0.056 | 0.022 | .894 |
| shawshank - citizenfour | -0.002 | -0.041 | 0.037 | .999 |
| prestige - citizenfour | -0.039 | -0.078 | 0.000 | .056 |
| summer - little miss sunshine | 0.023 | -0.008 | 0.054 | .306 |
| pulp fiction - little miss sunshine | 0.016 | -0.023 | 0.055 | .918 |
| shawshank - little miss sunshine | 0.030 | -0.009 | 0.070 | .257 |
| prestige - little miss sunshine | -0.006 | -0.045 | 0.033 | .999 |
| pulp fiction - summer | -0.007 | -0.046 | 0.032 | .999 |
| shawshank - summer | 0.007 | -0.032 | 0.046 | .999 |
| prestige - summer | -0.029 | -0.068 | 0.010 | .333 |
| shawshank - pulp fiction | 0.014 | -0.031 | 0.060 | .980 |
| pretige - pulp fiction | -0.022 | -0.068 | 0.024 | .842 |
| prestige - shawshank | -0.036 | -0.082 | 0.010 | .248 |
| **Gradient 5** | diff | lwr | upr | p adj |
| back to the future-12 years | 0.005 | -0.038 | 0.047 | .999 |
| citizenfour - 12 years | 0.043 | 0.007 | 0.079 | .008 |
| little miss sunshine - 12 years | -0.012 | -0.048 | 0.024 | .974 |
| summer - 12 years | -0.062 | -0.098 | -0.026 | <.001 |
| pulp fiction - 12 years | 0.006 | -0.036 | 0.049 | .999 |
| shawshank - 12 years | 0.029 | -0.014 | 0.071 | .441 |
| prestige - 12 years | -0.012 | -0.054 | 0.031 | .991 |
| citizenfour - backtothefuture | 0.038 | 0.002 | 0.074 | .031 |
| little miss sunshine - backtothefuture | -0.017 | -0.053 | 0.019 | .859 |
| summer - backtothefuture | -0.067 | -0.103 | -0.031 | <.001 |
| pulp fiction - backtothefuture | 0.002 | -0.041 | 0.044 | .999 |
| shawshank - backtothefuture | 0.024 | -0.018 | 0.067 | .671 |
| prestige - backtothefuture | -0.016 | -0.059 | 0.026 | .940 |
| little miss sunshine - citizenfour | -0.055 | -0.083 | -0.026 | <.001 |
| summer - citizenfour | -0.105 | -0.133 | -0.076 | <.001 |
| pulp fiction - citizenfour | -0.036 | -0.073 | 0.000 | .046 |
| shawshank - citizenfour | -0.014 | -0.050 | 0.022 | .942 |
| prestige - citizenfour | -0.054 | -0.091 | -0.018 | <.001 |
| summer - little miss sunshine | -0.050 | -0.079 | -0.022 | <.001 |
| pulp fiction - little miss sunshine | 0.018 | -0.018 | 0.054 | .792 |
| shawshank - little miss sunshine | 0.041 | 0.005 | 0.077 | .015 |
| prestige - little miss sunshine | 0.000 | -0.036 | 0.036 | .999 |
| pulp fiction - summer | 0.069 | 0.032 | 0.105 | <.001 |
| shawshank - summer | 0.091 | 0.055 | 0.127 | <.001 |
| prestige - summer | 0.051 | 0.014 | 0.087 | .001 |
| shawshank - pulp fiction | 0.023 | -0.020 | 0.065 | .743 |
| pretige - pulp fiction | -0.018 | -0.060 | 0.024 | .905 |
| prestige - shawshank | -0.041 | -0.083 | 0.002 | .073 |

*Note*. Tukey’s HSD output following ANOVA Model using Task_name (movie variable) to assess if a difference exists in Gradient scores across the movies. Significant results across pairwise comparisons indicate a difference in the average gradient score between the respective films.

**Supplementary Table 4 – Estimated Marginal Means (Gradient) Comparing to 0**

| Gradient | Movie | *emmean* | *SE* | *df* | *t* value | *p* value |
| --- | --- | --- | --- | --- | --- | --- |
| gradient_1 | 12 Years a Slave | -0.02 | 0.01 | 3733 | -1.75 | .080 |
| gradient_1 | Back to the Future | 0.02 | 0.01 | 3733 | 1.25 | .213 |
| gradient_1 | Citizenfour | 0.02 | 0.01 | 3733 | 2.02 | .044 |
| gradient_1 | Little Miss Sunshine | -0.07 | 0.01 | 3733 | -7.91 | <.001 |
| gradient_1 | 500 Days of Summer | -0.06 | 0.01 | 3733 | -6.50 | <.001 |
| gradient_1 | Pulp Fiction | -0.01 | 0.01 | 3733 | -1.01 | .311 |
| gradient_1 | Shawshank Redemption | 0.03 | 0.01 | 3733 | 2.16 | .031 |
| gradient_1 | The Prestige | 0.03 | 0.01 | 3733 | 2.28 | .023 |
| gradient_2 | 12 Years a Slave | -0.04 | 0.01 | 3733 | -2.71 | .007 |
| gradient_2 | Back to the Future | 0.00 | 0.01 | 3733 | -0.01 | .991 |
| gradient_2 | Citizenfour | -0.03 | 0.01 | 3733 | -3.00 | .003 |
| gradient_2 | Little Miss Sunshine | -0.12 | 0.01 | 3733 | -12.89 | <.001 |
| gradient_2 | 500 Days of Summer | 0.01 | 0.01 | 3733 | 1.06 | .288 |
| gradient_2 | Pulp Fiction | -0.02 | 0.01 | 3733 | -1.50 | .134 |
| gradient_2 | Shawshank Redemption | -0.03 | 0.01 | 3733 | -1.95 | .051 |
| gradient_2 | The Prestige | 0.03 | 0.01 | 3733 | 1.88 | .060 |
| gradient_3 | 12 Years a Slave | -0.02 | 0.01 | 3733 | -1.86 | .064 |
| gradient_3 | Back to the Future | -0.02 | 0.01 | 3733 | -1.62 | .105 |
| gradient_3 | Citizenfour | -0.05 | 0.01 | 3733 | -6.50 | <.001 |
| gradient_3 | Little Miss Sunshine | -0.04 | 0.01 | 3733 | -4.38 | <.001 |
| gradient_3 | 500 Days of Summer | 0.04 | 0.01 | 3733 | 4.77 | <.001 |
| gradient_3 | Pulp Fiction | 0.00 | 0.01 | 3733 | -0.03 | .975 |
| gradient_3 | Shawshank Redemption | -0.02 | 0.01 | 3733 | -1.77 | .078 |
| gradient_3 | The Prestige | 0.03 | 0.01 | 3733 | 2.40 | .016 |
| gradient_4 | 12 Years a Slave | 0.02 | 0.01 | 3733 | 1.83 | .068 |
| gradient_4 | Back to the Future | -0.02 | 0.01 | 3733 | -1.70 | .090 |
| gradient_4 | Citizenfour | 0.02 | 0.01 | 3733 | 2.74 | .006 |
| gradient_4 | Little Miss Sunshine | -0.01 | 0.01 | 3733 | -1.86 | .063 |
| gradient_4 | 500 Days of Summer | 0.01 | 0.01 | 3733 | 1.37 | .172 |
| gradient_4 | Pulp Fiction | 0.00 | 0.01 | 3733 | 0.25 | .799 |
| gradient_4 | Shawshank Redemption | 0.02 | 0.01 | 3733 | 1.61 | .109 |
| gradient_4 | The Prestige | -0.02 | 0.01 | 3733 | -1.77 | .077 |
| gradient_5 | 12 Years a Slave | 0.00 | 0.01 | 3733 | -0.43 | .671 |
| gradient_5 | Back to the Future | 0.00 | 0.01 | 3733 | 0.05 | .959 |
| gradient_5 | Citizenfour | 0.04 | 0.01 | 3733 | 5.80 | <.001 |
| gradient_5 | Little Miss Sunshine | -0.02 | 0.01 | 3733 | -2.43 | .015 |
| gradient_5 | 500 Days of Summer | -0.07 | 0.01 | 3733 | -9.96 | <.001 |
| gradient_5 | Pulp Fiction | 0.00 | 0.01 | 3733 | 0.21 | .834 |
| gradient_5 | Shawshank Redemption | 0.02 | 0.01 | 3733 | 2.49 | .013 |
| gradient_5 | The Prestige | -0.02 | 0.01 | 3733 | -1.61 | .108 |

*Note*. Estimated Marginal Means (EMMs) were computed for each model, as the ANOVA indicated a significant difference in the variability of gradient scores across films. For the purposes of our analysis, we computed a series of EMMs to assess if the gradient score associated with each film was significantly different from 0. The outcome of these tests suggests significant heterogeneity across the films in each gradient.

**Supplementary Table 5 – ANOVA (PCAs)**

| *Intrusive Distraction* (PCA_1) | *df* | Sum Sq | Mean Sq | *F* value | *p* value |
| --- | --- | --- | --- | --- | --- |
| Task_name | 7 | 144 | 20.544 | 21.33 | <.001 |
| Residuals | 3733 | 3596 | 0.963 |  |  |
| *Sensory Engagement* (PCA_2) |  |  |  |  |  |
| Task_name | 7 | 30 | 4.328 | 4.355 | <.001 |
| Residuals | 3733 | 3710 | 0.994 |  |  |
| *Episodic Knowledge* (PCA_3) |  |  |  |  |  |
| Task_name | 7 | 64 | 9.138 | 9.279 | <.001 |
| Residuals | 3733 | 3676 | 0.985 |  |  |
| *Deliberate Detail* (PCA_4) |  |  |  |  |  |
| Task_name | 7 | 16 | 2.2621 | 2.268 | .027 |
| Residuals | 3733 | 3724 | 0.9976 |  |  |
| *Positive Engagement* (PCA_5) |  |  |  |  |  |
| Task_name | 7 | 410 | 58.53 | 65.6 | <.001 |
| Residuals | 3733 | 3330 | 0.89 |  |  |

*Note.* ANOVA Model Outcome using Task_name (movie variable) to assess if a difference exists in thought component scores across the movies (PCA_1 = *Intrusive Distraction*, PCA_2 = *Sensory Engagement*, PCA_3 = *Episodic Knowledge*, PCA_4 = *Deliberate Detail*, PCA_5 = *Positive Engagement*) . Significant results for the main effect of Task_name suggest there is at least one difference in thought component scores in one of the films.

**Supplementary Table 6 – Tukey’s HSD Pairwise Comparisons (PCAs)**

| ***Intrusive Distraction* (PCA_1)** | diff | lwr | upr | p adj |
| --- | --- | --- | --- | --- |
| back to the future-12 years | 0.044 | -0.191 | 0.279 | .999 |
| citizenfour - 12 years | 0.583 | 0.383 | 0.783 | <.001 |
| little miss sunshine - 12 years | 0.197 | -0.003 | 0.397 | .057 |
| summer - 12 years | 0.130 | -0.070 | 0.331 | .501 |
| pulp fiction - 12 years | 0.321 | 0.086 | 0.556 | .001 |
| shawshank - 12 years | 0.032 | -0.203 | 0.267 | .999 |
| prestige - 12 years | 0.342 | 0.107 | 0.577 | <.001 |
| citizenfour - backtothefuture | 0.539 | 0.339 | 0.739 | <.001 |
| little miss sunshine - backtothefuture | 0.154 | -0.046 | 0.354 | .278 |
| summer - backtothefuture | 0.087 | -0.114 | 0.287 | .895 |
| pulp fiction - backtothefuture | 0.277 | 0.042 | 0.512 | .008 |
| shawshank - backtothefuture | -0.012 | -0.247 | 0.223 | .999 |
| prestige - backtothefuture | 0.298 | 0.063 | 0.533 | .003 |
| little miss sunshine - citizenfour | -0.385 | -0.543 | -0.228 | <.001 |
| summer - citizenfour | -0.452 | -0.610 | -0.294 | <.001 |
| pulp fiction - citizenfour | -0.262 | -0.462 | -0.062 | .002 |
| shawshank - citizenfour | -0.551 | -0.751 | -0.351 | <.001 |
| prestige - citizenfour | -0.241 | -0.441 | -0.041 | .006 |
| summer - little miss sunshine | -0.067 | -0.225 | 0.091 | .904 |
| pulp fiction - little miss sunshine | 0.124 | -0.076 | 0.324 | .568 |
| shawshank - little miss sunshine | -0.165 | -0.365 | 0.035 | .192 |
| prestige - little miss sunshine | 0.145 | -0.056 | 0.345 | .357 |
| pulp fiction - summer | 0.191 | -0.010 | 0.391 | .076 |
| shawshank - summer | -0.099 | -0.299 | 0.102 | .812 |
| prestige - summer | 0.211 | 0.011 | 0.412 | .030 |
| shawshank - pulp fiction | -0.289 | -0.524 | -0.054 | .005 |
| pretige - pulp fiction | 0.021 | -0.214 | 0.256 | .999 |
| prestige - shawshank | 0.310 | 0.075 | 0.545 | .002 |
| ***Sensory Engagement* (PCA_2)** | diff | lwr | upr | p adj |
| back to the future-12 years | -0.241 | -0.480 | -0.002 | .046 |
| citizenfour - 12 years | -0.231 | -0.434 | -0.028 | .013 |
| little miss sunshine - 12 years | -0.181 | -0.384 | 0.022 | .123 |
| summer - 12 years | -0.010 | -0.213 | 0.194 | .999 |
| pulp fiction - 12 years | -0.106 | -0.345 | 0.132 | .880 |
| shawshank - 12 years | -0.067 | -0.306 | 0.171 | .990 |
| prestige - 12 years | -0.104 | -0.343 | 0.134 | .890 |
| citizenfour - backtothefuture | 0.010 | -0.193 | 0.213 | .999 |
| little miss sunshine - backtothefuture | 0.060 | -0.143 | 0.263 | .986 |
| summer - backtothefuture | 0.231 | 0.028 | 0.435 | .013 |
| pulp fiction - backtothefuture | 0.135 | -0.104 | 0.373 | .679 |
| shawshank - backtothefuture | 0.174 | -0.065 | 0.412 | .348 |
| prestige - backtothefuture | 0.137 | -0.102 | 0.375 | .663 |
| little miss sunshine - citizenfour | 0.050 | -0.110 | 0.210 | .981 |
| summer - citizenfour | 0.221 | 0.061 | 0.382 | .001 |
| pulp fiction - citizenfour | 0.125 | -0.079 | 0.328 | .579 |
| shawshank - citizenfour | 0.163 | -0.040 | 0.367 | .222 |
| prestige - citizenfour | 0.127 | -0.077 | 0.330 | .559 |
| summer - little miss sunshine | 0.171 | 0.011 | 0.332 | .027 |
| pulp fiction - little miss sunshine | 0.075 | -0.128 | 0.278 | .954 |
| shawshank - little miss sunshine | 0.114 | -0.090 | 0.317 | .690 |
| prestige - little miss sunshine | 0.077 | -0.127 | 0.280 | .947 |
| pulp fiction - summer | -0.097 | -0.300 | 0.107 | .839 |
| shawshank - summer | -0.058 | -0.261 | 0.146 | .989 |
| prestige - summer | -0.095 | -0.298 | 0.109 | .852 |
| shawshank - pulp fiction | 0.039 | -0.200 | 0.278 | .999 |
| pretige - pulp fiction | 0.002 | -0.237 | 0.241 | .999 |
| prestige - shawshank | -0.037 | -0.276 | 0.202 | .999 |
| ***Episodic Knowledge* (PCA_3)** | diff | lwr | upr | p adj |
| back to the future-12 years | 0.061 | -0.177 | 0.298 | .994 |
| citizenfour - 12 years | -0.230 | -0.432 | -0.027 | .013 |
| little miss sunshine - 12 years | -0.125 | -0.327 | 0.077 | .570 |
| summer - 12 years | 0.024 | -0.179 | 0.226 | .999 |
| pulp fiction - 12 years | -0.391 | -0.629 | -0.154 | <.001 |
| shawshank - 12 years | -0.085 | -0.322 | 0.153 | .961 |
| prestige - 12 years | -0.192 | -0.430 | 0.045 | .217 |
| citizenfour - backtothefuture | -0.290 | -0.493 | -0.088 | <.001 |
| little miss sunshine - backtothefuture | -0.186 | -0.388 | 0.017 | .100 |
| summer - backtothefuture | -0.037 | -0.240 | 0.165 | .999 |
| pulp fiction - backtothefuture | -0.452 | -0.690 | -0.215 | <.001 |
| shawshank - backtothefuture | -0.145 | -0.383 | 0.092 | .582 |
| prestige - backtothefuture | -0.253 | -0.490 | -0.015 | .028 |
| little miss sunshine - citizenfour | 0.105 | -0.054 | 0.264 | .485 |
| summer - citizenfour | 0.253 | 0.094 | 0.413 | <.001 |
| pulp fiction - citizenfour | -0.162 | -0.364 | 0.040 | .229 |
| shawshank - citizenfour | 0.145 | -0.057 | 0.347 | .367 |
| prestige - citizenfour | 0.038 | -0.165 | 0.240 | .999 |
| summer - little miss sunshine | 0.149 | -0.011 | 0.308 | .090 |
| pulp fiction - little miss sunshine | -0.267 | -0.469 | -0.064 | .002 |
| shawshank - little miss sunshine | 0.040 | -0.162 | 0.242 | .999 |
| prestige - little miss sunshine | -0.067 | -0.269 | 0.135 | .973 |
| pulp fiction - summer | -0.415 | -0.618 | -0.213 | <.001 |
| shawshank - summer | -0.108 | -0.311 | 0.094 | .737 |
| prestige - summer | -0.216 | -0.418 | -0.013 | .027 |
| shawshank - pulp fiction | 0.307 | 0.069 | 0.544 | .002 |
| pretige - pulp fiction | 0.199 | -0.038 | 0.437 | .177 |
| prestige - shawshank | -0.107 | -0.345 | 0.130 | .870 |
| ***Deliberate Detail* (PCA_4)** | diff | lwr | upr | p adj |
| back to the future-12 years | 0.112 | -0.127 | 0.351 | .849 |
| citizenfour - 12 years | 0.132 | -0.072 | 0.335 | .506 |
| little miss sunshine - 12 years | 0.007 | -0.196 | 0.211 | .999 |
| summer - 12 years | 0.032 | -0.172 | 0.236 | .999 |
| pulp fiction - 12 years | 0.109 | -0.130 | 0.348 | .865 |
| shawshank - 12 years | 0.206 | -0.033 | 0.445 | .150 |
| prestige - 12 years | 0.024 | -0.215 | 0.263 | .999 |
| citizenfour - backtothefuture | 0.020 | -0.184 | 0.223 | .999 |
| little miss sunshine - backtothefuture | -0.105 | -0.308 | 0.099 | .775 |
| summer - backtothefuture | -0.080 | -0.284 | 0.124 | .935 |
| pulp fiction - backtothefuture | -0.003 | -0.242 | 0.236 | .999 |
| shawshank - backtothefuture | 0.094 | -0.145 | 0.334 | .933 |
| prestige - backtothefuture | -0.088 | -0.327 | 0.151 | .953 |
| little miss sunshine - citizenfour | -0.125 | -0.285 | 0.036 | .263 |
| summer - citizenfour | -0.100 | -0.260 | 0.061 | .562 |
| pulp fiction - citizenfour | -0.023 | -0.226 | 0.181 | .999 |
| shawshank - citizenfour | 0.074 | -0.129 | 0.278 | .955 |
| prestige - citizenfour | -0.108 | -0.312 | 0.096 | .745 |
| summer - little miss sunshine | 0.025 | -0.136 | 0.185 | .999 |
| pulp fiction - little miss sunshine | 0.102 | -0.102 | 0.305 | .798 |
| shawshank - little miss sunshine | 0.199 | -0.004 | 0.403 | .061 |
| prestige - little miss sunshine | 0.017 | -0.187 | 0.220 | .999 |
| pulp fiction - summer | 0.077 | -0.127 | 0.281 | .946 |
| shawshank - summer | 0.174 | -0.030 | 0.378 | .158 |
| prestige - summer | -0.008 | -0.212 | 0.196 | .999 |
| shawshank - pulp fiction | 0.097 | -0.142 | 0.336 | .922 |
| pretige - pulp fiction | -0.085 | -0.324 | 0.154 | .961 |
| prestige - shawshank | -0.182 | -0.422 | 0.057 | .286 |
| ***Positive Engagement* (PCA_5)** | diff | lwr | upr | *p*-val |
| back to the future-12 years | 1.151 | 0.925 | 1.377 | <.001 |
| citizenfour - 12 years | 0.453 | 0.260 | 0.645 | <.001 |
| little miss sunshine - 12 years | 0.832 | 0.640 | 1.025 | <.001 |
| summer - 12 years | 1.078 | 0.885 | 1.270 | <.001 |
| pulp fiction - 12 years | 0.860 | 0.634 | 1.086 | <.001 |
| shawshank - 12 years | 0.953 | 0.727 | 1.179 | <.001 |
| prestige - 12 years | 0.524 | 0.298 | 0.750 | <.001 |
| citizenfour - backtothefuture | -0.698 | -0.891 | -0.506 | <.001 |
| little miss sunshine - backtothefuture | -0.319 | -0.511 | -0.126 | <.001 |
| summer - backtothefuture | -0.074 | -0.266 | 0.119 | .944 |
| pulp fiction - backtothefuture | -0.291 | -0.517 | -0.065 | .002 |
| shawshank - backtothefuture | -0.198 | -0.424 | 0.028 | .136 |
| prestige - backtothefuture | -0.627 | -0.854 | -0.401 | <.001 |
| little miss sunshine - citizenfour | 0.380 | 0.228 | 0.531 | <.001 |
| summer - citizenfour | 0.625 | 0.473 | 0.777 | <.001 |
| pulp fiction - citizenfour | 0.407 | 0.215 | 0.600 | <.001 |
| shawshank - citizenfour | 0.500 | 0.308 | 0.693 | <.001 |
| prestige - citizenfour | 0.071 | -0.122 | 0.263 | .953 |
| summer - little miss sunshine | 0.245 | 0.093 | 0.397 | <.001 |
| pulp fiction - little miss sunshine | 0.027 | -0.165 | 0.220 | 1.000 |
| shawshank - little miss sunshine | 0.121 | -0.072 | 0.313 | .552 |
| prestige - little miss sunshine | -0.309 | -0.501 | -0.116 | <.001 |
| pulp fiction - summer | -0.218 | -0.411 | -0.025 | .014 |
| shawshank - summer | -0.125 | -0.317 | 0.068 | .508 |
| prestige - summer | -0.554 | -0.747 | -0.361 | <.001 |
| shawshank - pulp fiction | 0.093 | -0.133 | 0.319 | .917 |
| pretige - pulp fiction | -0.336 | -0.562 | -0.110 | <.001 |
| prestige - shawshank | -0.429 | -0.655 | -0.203 | <.001 |

*Note*. Tukey’s HSD output following ANOVA Model using Task_name (movie variable) to assess if a difference exists in thought component (PCA) scores across the movies. Significant results across pairwise comparisons indicate a difference in the average thought component score between the respective films.

**Supplementary Table 7 – Estimated Marginal Means (PCAs) Comparing to 0**

| Thought Component | Movie | *emmean* | *SE* | *df* | *t* value | *p* value |
| --- | --- | --- | --- | --- | --- | --- |
| **PCA 1** |  |  |  |  |  |  |
| *Intrusive Distraction* | 12 Years a Slave | -0.24 | 0.05 | 3733 | -4.32 | <.001 |
| *Intrusive Distraction* | Back to the Future | -0.19 | 0.05 | 3733 | -3.53 | <.001 |
| *Intrusive Distraction* | Citizenfour | 0.35 | 0.04 | 3733 | 9.41 | <.001 |
| *Intrusive Distraction* | Little Miss Sunshine | -0.04 | 0.04 | 3733 | -1.08 | .281 |
| *Intrusive Distraction* | 500 Days of Summer | -0.11 | 0.04 | 3733 | -2.89 | .004 |
| *Intrusive Distraction* | Pulp Fiction | 0.08 | 0.05 | 3733 | 1.54 | .125 |
| *Intrusive Distraction* | Shawshank Redemption | -0.21 | 0.05 | 3733 | -3.74 | <.001 |
| *Intrusive Distraction* | The Prestige | 0.10 | 0.05 | 3733 | 1.92 | .056 |
| **PCA 2** |  |  |  |  |  |  |
| *Sensory Engagement* | 12 Years a Slave | 0.12 | 0.06 | 3733 | 2.24 | .025 |
| *Sensory Engagement* | Back to the Future | -0.12 | 0.06 | 3733 | -2.09 | .037 |
| *Sensory Engagement* | Citizenfour | -0.11 | 0.04 | 3733 | -2.84 | .005 |
| *Sensory Engagement* | Little Miss Sunshine | -0.06 | 0.04 | 3733 | -1.50 | .134 |
| *Sensory Engagement* | 500 Days of Summer | 0.12 | 0.04 | 3733 | 3.08 | .002 |
| *Sensory Engagement* | Pulp Fiction | 0.02 | 0.06 | 3733 | 0.34 | .737 |
| *Sensory Engagement* | Shawshank Redemption | 0.06 | 0.06 | 3733 | 1.04 | .301 |
| *Sensory Engagement* | The Prestige | 0.02 | 0.06 | 3733 | 0.37 | .711 |
| **PCA 3** |  |  |  |  |  |  |
| *Episodic Knowledge* | 12 Years a Slave | 0.12 | 0.06 | 3733 | 2.08 | .037 |
| *Episodic Knowledge* | Back to the Future | 0.18 | 0.06 | 3733 | 3.18 | .002 |
| *Episodic Knowledge* | Citizenfour | -0.11 | 0.04 | 3733 | -3.08 | .002 |
| *Episodic Knowledge* | Little Miss Sunshine | -0.01 | 0.04 | 3733 | -0.26 | .797 |
| *Episodic Knowledge* | 500 Days of Summer | 0.14 | 0.04 | 3733 | 3.73 | <.001 |
| *Episodic Knowledge* | Pulp Fiction | -0.28 | 0.06 | 3733 | -4.99 | <.001 |
| *Episodic Knowledge* | Shawshank Redemption | 0.03 | 0.06 | 3733 | 0.55 | .580 |
| *Episodic Knowledge* | The Prestige | -0.08 | 0.06 | 3733 | -1.39 | .166 |
| **PCA 4** |  |  |  |  |  |  |
| *Deliberate Detail* | 12 Years a Slave | -0.07 | 0.06 | 3733 | -1.28 | .201 |
| *Deliberate Detail* | Back to the Future | 0.04 | 0.06 | 3733 | 0.73 | .467 |
| *Deliberate Detail* | Citizenfour | 0.06 | 0.04 | 3733 | 1.62 | .106 |
| *Deliberate Detail* | Little Miss Sunshine | -0.06 | 0.04 | 3733 | -1.71 | .087 |
| *Deliberate Detail* | 500 Days of Summer | -0.04 | 0.04 | 3733 | -1.05 | .295 |
| *Deliberate Detail* | Pulp Fiction | 0.04 | 0.06 | 3733 | 0.68 | .498 |
| *Deliberate Detail* | Shawshank Redemption | 0.14 | 0.06 | 3733 | 2.42 | .016 |
| *Deliberate Detail* | The Prestige | -0.05 | 0.06 | 3733 | -0.85 | .395 |
| **PCA 5** |  |  |  |  |  |  |
| *Positive Engagement* | 12 Years a Slave | -0.75 | 0.05 | 3733 | -14.20 | <.001 |
| *Positive Engagement* | Back to the Future | 0.40 | 0.05 | 3733 | 7.64 | <.001 |
| *Positive Engagement* | Citizenfour | -0.30 | 0.04 | 3733 | -8.37 | <.001 |
| *Positive Engagement* | Little Miss Sunshine | 0.08 | 0.04 | 3733 | 2.37 | .018 |
| *Positive Engagement* | 500 Days of Summer | 0.33 | 0.04 | 3733 | 9.27 | <.001 |
| *Positive Engagement* | Pulp Fiction | 0.11 | 0.05 | 3733 | 2.11 | .035 |
| *Positive Engagement* | Shawshank Redemption | 0.20 | 0.05 | 3733 | 3.88 | <.001 |
| *Positive Engagement* | The Prestige | -0.22 | 0.05 | 3733 | -4.26 | <.001 |

*Note.* Estimated Marginal Means (EMMs) were computed for each model, as the ANOVA indicated a significant difference in the variability of thought component scores across films. For the purposes of our analysis, we computed a series of EMMs to assess if the thought score associated with each film was significantly different from 0. The outcome of these tests suggests significant heterogeneity across the films in each thought component.

**Supplementary Table 8 – LMM Thought & Brain Association (Subject ID Random)**

| Subject ID Random Effects |  |  |  |  |  |
| --- | --- | --- | --- | --- | --- |
| ***Intrusive Distraction* (PCA_1)** | Estimate | *SE* | *df* | *t-val* | *p-*val |
| (Intercept) | -.01 | 0.04 | 226 | -0.17 | .864 |
| gradient_1 | .00 | 0.07 | 3563 | 0.06 | .949 |
| gradient_2 | .09 | 0.06 | 3565 | 1.34 | .181 |
| gradient_3 | -.23 | 0.08 | 3551 | -2.89 | .004 |
| gradient_4 | -.10 | 0.09 | 3561 | -1.07 | .283 |
| gradient_5 | .12 | 0.09 | 3543 | 1.33 | .183 |
| ***Sensory Engagement* (PCA_2)** | Estimate | *SE* | *df* | *t-val* | *p-*val |
| (Intercept) | .00 | 0.05 | 225 | 0.05 | .962 |
| gradient_1 | -.13 | 0.06 | 3543 | -2.03 | .042 |
| gradient_2 | .04 | 0.06 | 3544 | 0.72 | .473 |
| gradient_3 | .09 | 0.07 | 3536 | 1.29 | .199 |
| gradient_4 | .13 | 0.08 | 3542 | 1.62 | .105 |
| gradient_5 | .19 | 0.08 | 3530 | 2.22 | .026 |
| ***Episodic Knowledge* (PCA_3)** | Estimate | *SE* | *df* | *t-val* | *p-*val |
| (Intercept) | .00 | 0.04 | 226 | 0.05 | .960 |
| gradient_1 | .01 | 0.07 | 3576 | 0.19 | .851 |
| gradient_2 | .05 | 0.07 | 3577 | 0.80 | .423 |
| gradient_3 | .06 | 0.08 | 3561 | 0.74 | .459 |
| gradient_4 | .16 | 0.10 | 3573 | 1.69 | .091 |
| gradient_5 | .14 | 0.10 | 3551 | 1.45 | .148 |
| ***Deliberate Detail* (PCA_4)** | Estimate | *SE* | *df* | *t-val* | *p-*val |
| (Intercept) | .00 | 0.04 | 225 | 0.06 | .950 |
| gradient_1 | .01 | 0.07 | 3572 | 0.07 | .941 |
| gradient_2 | .09 | 0.07 | 3574 | 1.34 | .180 |
| gradient_3 | -.02 | 0.08 | 3559 | -0.29 | .772 |
| gradient_4 | -.01 | 0.10 | 3570 | -0.14 | .889 |
| gradient_5 | .02 | 0.10 | 3548 | 0.18 | .854 |
| ***Positive Engagement* (PCA_5)** | Estimate | *SE* | *df* | *t-val* | *p-*val |
| (Intercept) | -.01 | 0.03 | 228 | -0.20 | .841 |
| gradient_1 | .03 | 0.08 | 3625 | 0.37 | .712 |
| gradient_2 | -.14 | 0.07 | 3628 | -1.90 | .057 |
| gradient_3 | .16 | 0.09 | 3602 | 1.78 | .075 |
| gradient_4 | .14 | 0.10 | 3621 | 1.37 | .172 |
| gradient_5 | -.08 | 0.10 | 3584 | -0.79 | .429 |

Note. Linear Mixed Model predicting thought patterns using gradient maps 1-5, including participant as a random intercept effect. *P-*values adjusted using FDR.

**Supplementary Table 9 – LMM Thought & Brain Association (Subject ID + Movie Random)**

| Subject ID + Movie Random Effects |  |  |  |  |  |
| --- | --- | --- | --- | --- | --- |
| ***Intrusive Distraction* (PCA_1)** | Estimate | *SE* | *df* | *t-val* | *p-*val |
| (Intercept) | -.03 | 0.08 | 12 | -0.34 | .740 |
| gradient_1 | -.02 | 0.07 | 3555 | -0.30 | .764 |
| gradient_2 | .02 | 0.06 | 3558 | 0.35 | .726 |
| gradient_3 | -.13 | 0.08 | 3549 | -1.61 | .107 |
| gradient_4 | -.01 | 0.09 | 3557 | -0.10 | .919 |
| gradient_5 | -.02 | 0.09 | 3539 | -0.17 | .869 |
| ***Sensory Engagement* (PCA_2)** | Estimate | *SE* | *df* | *t-val* | *p-*val |
| (Intercept) | .00 | 0.06 | 43 | 0.08 | .935 |
| gradient_1 | -.12 | 0.06 | 3540 | -1.90 | .057 |
| gradient_2 | .05 | 0.06 | 3544 | 0.93 | .350 |
| gradient_3 | .01 | 0.07 | 3533 | 0.20 | .844 |
| gradient_4 | .05 | 0.08 | 3541 | 0.65 | .518 |
| gradient_5 | .26 | 0.08 | 3529 | 3.10 | .002 |
| ***Episodic Knowledge* (PCA_3)** | Estimate | *SE* | *df* | *t-val* | *p-*val |
| (Intercept) | .00 | 0.07 | 14 | 0.02 | .988 |
| gradient_1 | .02 | 0.07 | 3569 | 0.26 | .795 |
| gradient_2 | .07 | 0.07 | 3573 | 0.98 | .326 |
| gradient_3 | .01 | 0.08 | 3561 | 0.10 | .920 |
| gradient_4 | .11 | 0.10 | 3571 | 1.16 | .246 |
| gradient_5 | .23 | 0.10 | 3549 | 2.35 | .019 |
| ***Deliberate Detail* (PCA_4)** | Estimate | *SE* | *df* | *t-val* | *p-*val |
| (Intercept) | .00 | 0.05 | 42 | 0.07 | .945 |
| gradient_1 | .00 | 0.07 | 3571 | -0.03 | .979 |
| gradient_2 | .07 | 0.07 | 3559 | 1.10 | .273 |
| gradient_3 | .00 | 0.08 | 3484 | 0.05 | .957 |
| gradient_4 | .01 | 0.10 | 3535 | 0.06 | .952 |
| gradient_5 | -.02 | 0.10 | 3499 | -0.21 | .832 |
| ***Positive Engagement* (PCA_5)** | Estimate | *SE* | *df* | *t-val* | *p-*val |
| (Intercept) | -.02 | 0.14 | 8 | -0.13 | .897 |
| gradient_1 | .04 | 0.07 | 3604 | 0.48 | .634 |
| gradient_2 | -.11 | 0.07 | 3608 | -1.64 | .101 |
| gradient_3 | .03 | 0.09 | 3588 | 0.35 | .723 |
| gradient_4 | .07 | 0.10 | 3603 | 0.69 | .492 |
| gradient_5 | .09 | 0.10 | 3570 | 0.92 | .360 |

Note. Linear Mixed Model predicting thought patterns using gradient maps 1-5, with participant and movie identity as random intercepts. *P-*values adjusted using FDR.

**Supplementary Table 10 - LMM Comprehension**

|  | Estimate | *SE* | *df* | *t*-value | *p*-val |
| --- | --- | --- | --- | --- | --- |
| (Intercept) | 2.987 | .039 | 226 | 77.484 | <.001 |
| *Intrusive Distraction* | -0.166 | .015 | 3735 | -10.864 | <.001 |
| *Sensory Engagement* | 0.016 | .017 | 3652 | 0.934 | .350 |
| *Episodic Knowledge* | 0.050 | .015 | 3728 | 3.358 | .001 |
| *Deliberate Detail* | -0.023 | .015 | 3732 | -1.56 | .149 |
| *Positive Engagement* | 0.119 | .014 | 3660 | 8.666 | <.001 |

Note. Linear Mixed Model predicting comprehension scores using gradient maps 1-5, with participants as random intercepts. *P-*values adjusted using FDR

**Supplementary Table 11 – Movie Comprehension Questions**

| Movie | Question | Answer |
| --- | --- | --- |
| The Prestige | What does Sarah say she wants from him? | Openness |
| The Prestige | What is the name of the trick the magician plans to base his show around? | Transported man |
| The Prestige | What playing card does the man spot under his glass? | King |
| The Prestige | What is said to be the truth about him? | He never loved Sarah |
| Pulp Fiction | What is item is originally said to be in the case? | Laundry |
| Pulp Fiction | What name does the male robber call the female robber? | Honey Bunny |
| Pulp Fiction | What is the female robbers real name? | Yolanda |
| Pulp Fiction | How much money does the woman give him? | $15,000 |
| The Shawshank Redemption | What tier did the guard say a man was missing? | Tier 2 |
| The Shawshank Redemption | What does the warden refer to the girl on the poster as? | Cupcake on the wall |
| The Shawshank Redemption | How many years did it take Andy to tunnel through the wall with the rock hammer? | Less than 20 |
| The Shawshank Redemption | What was Andy initially carving into his prison cell wall? | His initial ("A") |
| Back to the future | What year is the date on the newspaper? | 1955 |
| Back to the future | How much is the cup of coffee advertised for in the diner? | 5 cents |
| Back to the future | What drink does he ask for? | Pepsi |
| Back to the future | What does Marty call his dad when he catches him on the tree? | Peeping Tom |
| 12 Years a Slave | What is the lady making out of a corn husk? | Dolls |
| 12 Years a Slave | What instrument is the man playing? | Violin |
| 12 Years a Slave | What does the woman throw at the girl while she's dancing? | Glass bottle |
| 12 Years a Slave | Which item is on the list given to the man? | Places to visit |
| Citizenfour | What animal is seen on screen in the first scene? | Dog |
| Citizenfour | What president is being discussed in the phone call with David? | Barack Obama |
| Citizenfour | After 4 years in the military, who directly entered the NSA with 37 years of combined service? | William |
| Citizenfour | According to the movie, what historical event triggered the government to begin spying on citizens? | 9/11 |
| Little Miss Sunshine | What breakfast item did Olive order a la mode? | Waffles |
| Little Miss Sunshine | What colour is the Volkswagen Transporter that the family drives? | Yellow |
| Little Miss Sunshine | How does the family get into their car after taking it to the repair shop? | Pushing it |
| Little Miss Sunshine | What type of language does Richard claim is “the refuge of losers”? | Sarcasm |
| 500 Days of Summer | What event are both Summer and Tom attending? | Wedding |
| 500 Days of Summer | What piece of jewelry is Summer showing her friend when Tom abruptly leaves her party? | Engagement ring |
| 500 Days of Summer | On day 440, what time does Tom’s alarm clock go off? | 7AM |
| 500 Days of Summer | What book is Tom reading on the train, which he also gifts to Summer at her party? | Architecture of happiness |

**Supplementary Table 11 – multi-dimensional experience sampling (mDES)**

| **Dimension** | **Question** | **Scale Low (0)** | **Scale High (10)** |
| --- | --- | --- | --- |
| Focus | My thoughts were focused on an external task or activity | Not at all | Completely |
| Future | My thoughts involved future events: | Not at all | Completely |
| Past | My thoughts involved past events: | Not at all | Completely |
| Self | My thoughts involved myself: | Not at all | Completely |
| Other | My thoughts involved other people: | Not at all | Completely |
| Emotion | The emotion of my thoughts was: | Negative | Positive |
| Images | My thoughts were in the form of images | Not at all | Completely |
| Sounds | My thoughts were in the form of sounds | Not at all | Completely |
| Words | My thoughts were in the form of words | Not at all | Completely |
| Detailed | My thoughts were detailed and specific: | Not at all | Completely |
| Deliberate | My thoughts were: | Spontaneous | Deliberate |
| Problem | I was thinking about solutions to problems (or goals): | Not at all | Completely |
| Intrusive | My thoughts were intrusive: | Not at all | Completely |
| Knowledge | My thoughts contained information I already knew (e.g., knowledge or memories) | Not at all | Completely |
| Absorption | I was absorbed in the contents of my thoughts | Not at all | Completely |
| Distraction | My thoughts were distracting me from what I am doing | Not at all | Completely |

*Note.* List of 16-item multi-dimensional experience sampling (mDES) questions delivered during each probe in the experience sampling paradigm. Participants scores are recorded on a continuous scale from 0 (Not at all) to 10 (Completely).

**Supplementary Figure 1**

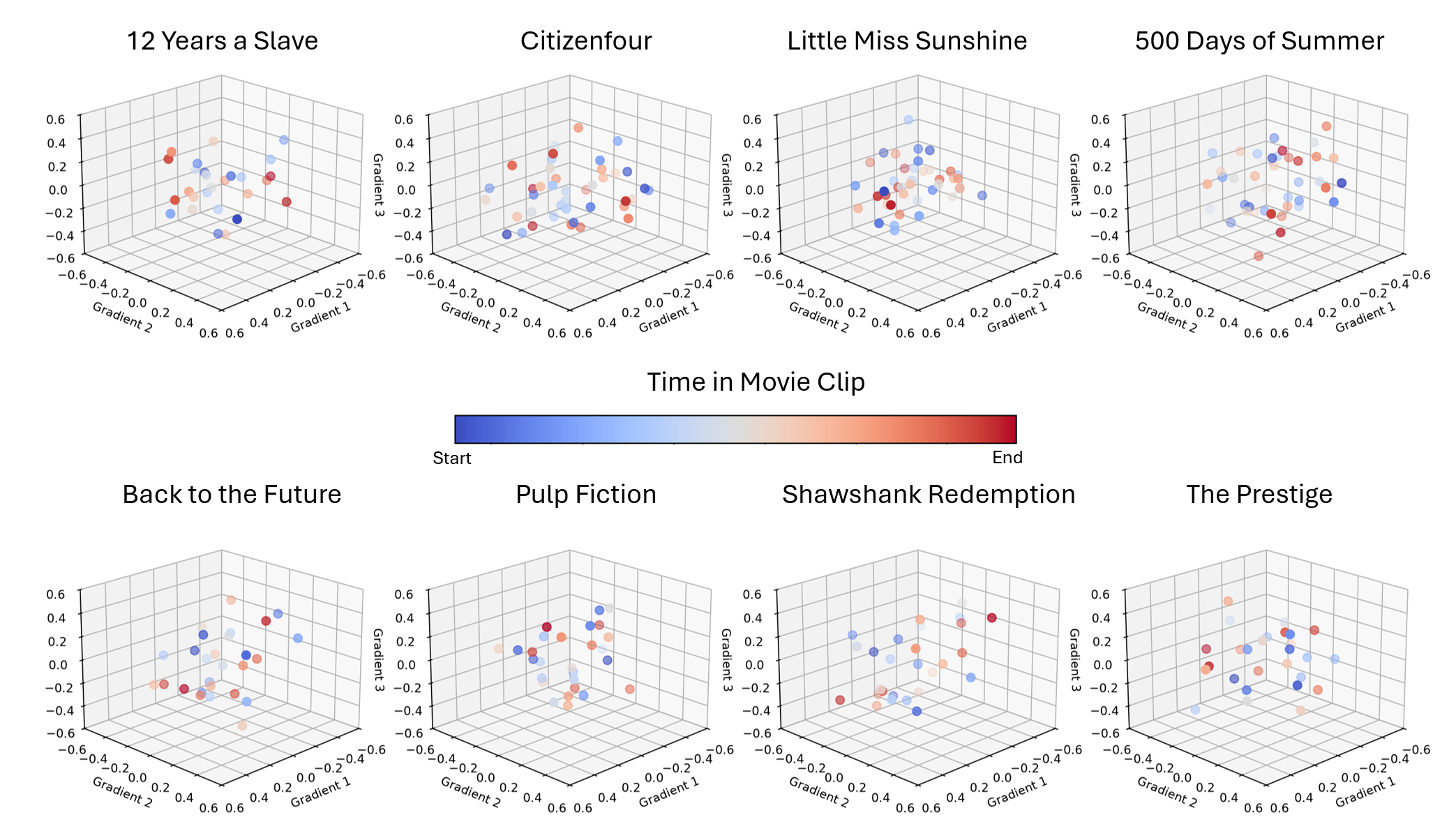

*Note.* Timecourse 3D scatter plot of ‘brain space’ trajectory across gradient coordinates using gradient 1, gradient 2, and gradient 3 for each film. Cooler points in blue are associated with earlier in the film clip, whereas warmer points in red are associated with the end of the clip.

**Supplementary Figure 2**

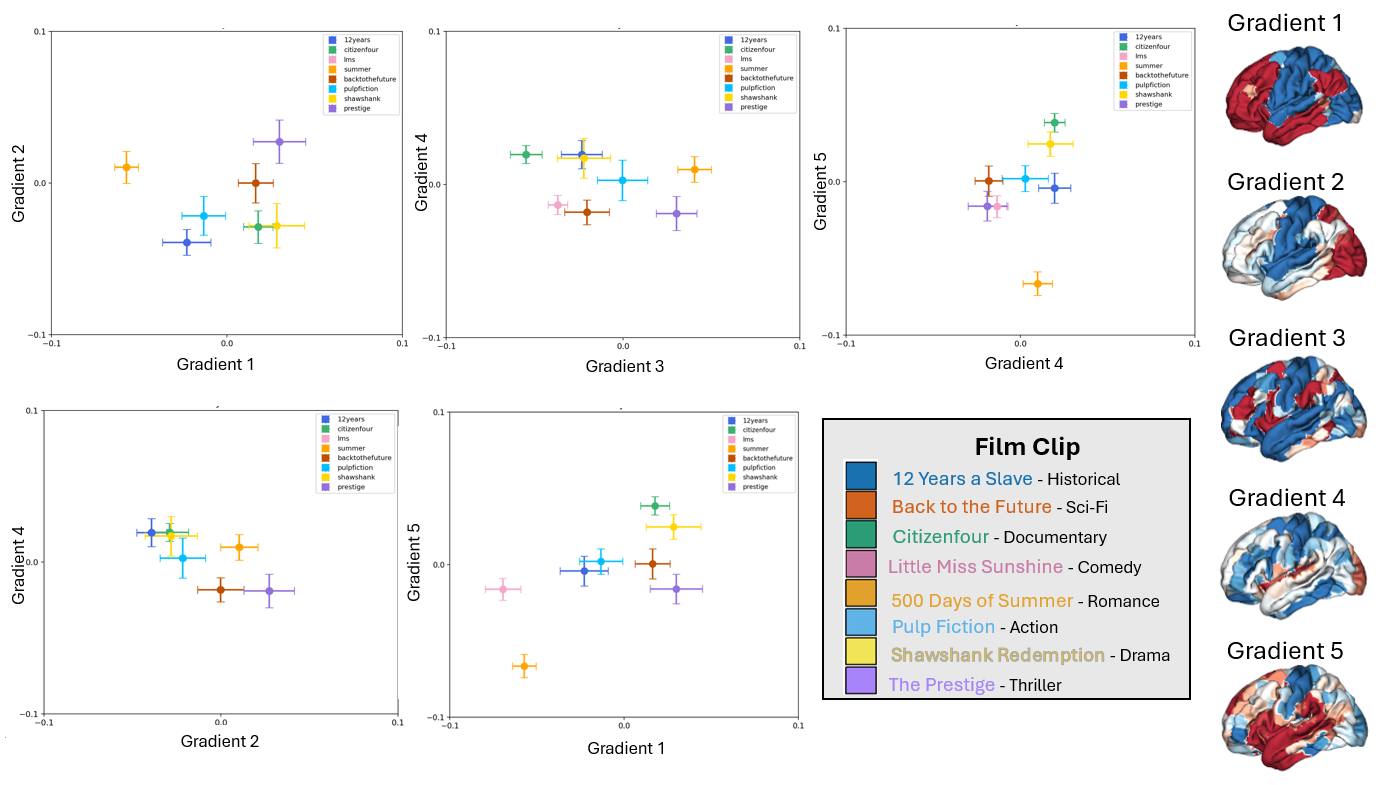

*Note.* Gradient scatter plots with 99% CI error bars illustrating the group-average association of each gradient combination with the eight films. The corresponding gradient maps are on the right.

**Supplementary Figure 3**

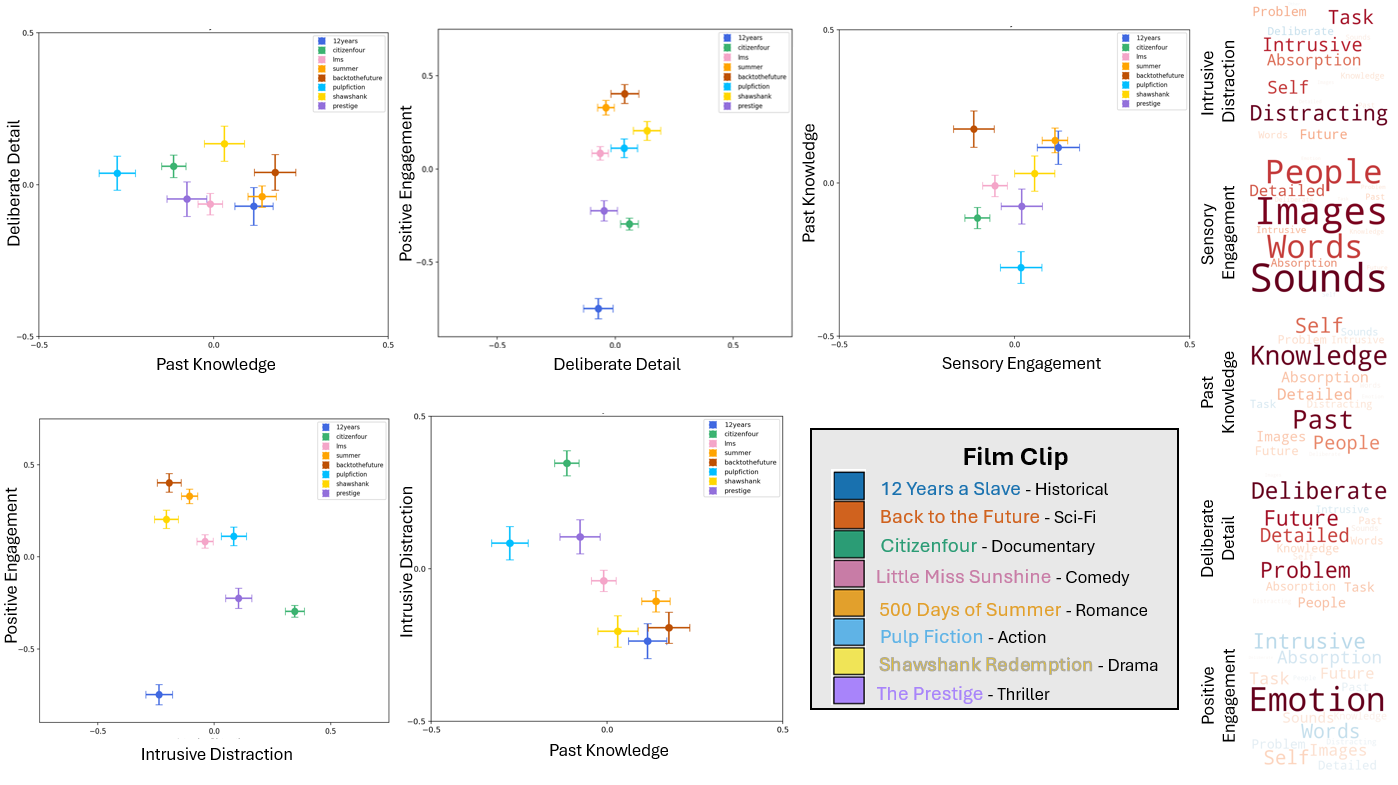

*Note.* Thought pattern scatter plots with 99% CI error bars illustrating the group-average association of each thought pattern combination with the eight films. The corresponding thought pattern word clouds are on the right.

**Supplementary Figure 4**

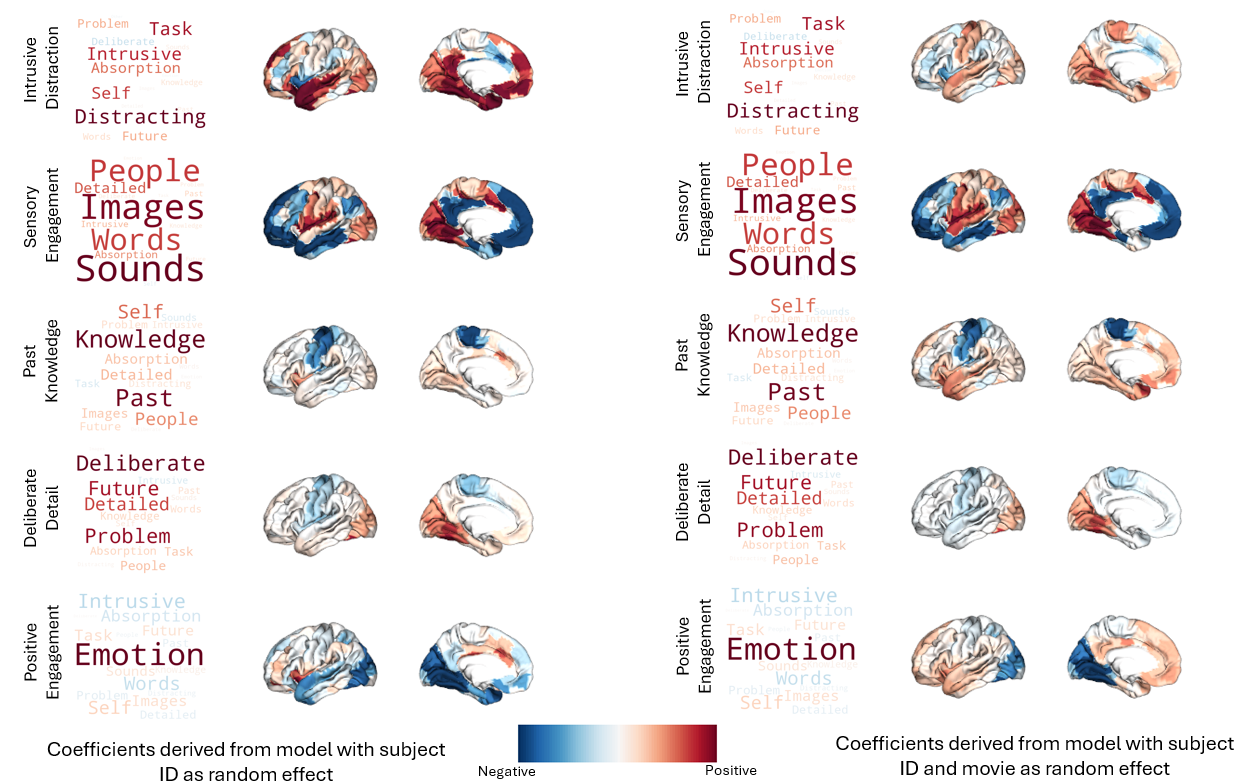

*Note.* Dot-product maps for each thought pattern in Set 1 and Set 2 linear mixed models. Brain plots are comprised of the gradient estimate effect and summarized into the plots on the right to the corresponding thought pattern. The colour bar indicates the directionality of the activation association (red = positive, blue = negative).
